# New specimens of *Bunaia woodwardi*, Clarke 1919 (Euchelicerata): A new member of Offacolidae providing insight supporting the Arachnomorpha

**DOI:** 10.1101/2024.03.18.585490

**Authors:** Lorenzo Lustri, Jonathan B. Antcliffe, Pierre Gueriau, Allison C. Daley

**Affiliations:** Institute of Earth Sciences, University of Lausanne, Géopolis, CH-1015 Lausanne, Switzerland; Université Paris-Saclay, CNRS, ministère de la Culture, UVSQ, MNHN, Institut photonique d’analyse non-destructive européen des matériaux anciens, 91192, Saint-Aubin, France

**Keywords:** Offacolidae, Euchelicerata, Vicissicaudata, Habeliida, Artiopoda, Arachnomorpha

## Abstract

The rapid early diversification of arthropods has made understanding internal relationships within the group fiendish. Particularly unresolved is the origin of Euchelicerata, a clade consisting of the Prosomapoda (comprising the extant Xiphosura and Arachnida and the extinct Chasmataspidida, Eurypterida and synziphosurines) and the extinct Offacolidae. Here we describe new material of the Silurian ‘synziphosurine’ *Bunaia woodwardi* that reveals previously unknown features of its ventral anatomy: a pair of elongated chelicerae in the prosoma, followed posteriorly by five pairs of biramous appendages, a first pre-abdomen somite bearing a pair of paddle-like uniramous appendages (exopods), and a ventral pretelsonic process. Phylogenetic analyses retrieve *B. woodwardi* as an Offacolidae closely related to *Setapedites abundantis* from the early Ordovician Fezouata Biota. An anatomical comparison of the pretelsonic process of *B. woodwardi*, also present in *Setapedites*, with the posterior trunk morphologies of other Offacolidae, Habeliida and Vicissicaudata, suggests a possible homologous appendicular origin. This proposed apomorphic character supports a monophyletic Arachnomorpha, formed of Vicissicaudata, Habeliida and Euchelicerata. The establishment of this new homology could help to clarify the highly enigmatic phylogeny at the base of the euchelicerates as well as the sequence of character acquisition during their early evolution.

## 1. Introduction

Arthropoda is divided into several major groupings whose relationships are highly controversial and subject to rapid revision and terminological changes [1–7]. One such major division within the arthropods is Euchelicerata, a highly diverse clade that includes spiders, scorpions, and mites (arachnids) as well as horseshoe crabs (xiphosurans). Euchelicerata also contains several important extinct forms such as sea scorpions (eurypterids), chasmataspidids, and synziphosurines. The enigmatic synziphosurines, which superficially resemble modern horseshoe crabs, dwelt in the seas from the Early Ordovician (Tremadocian (ca. 484 million years ago (Ma)) [8] to the Late Carboniferous (Mississippian (ca. 323 Ma)) [9]. They scratched out a living searching for food on the sediment surface of these ancient seabeds and may in some species have had a limited swimming ability, rather like modern horseshoe crabs.

Though the phylogenetic position of these synziphosurines has proven problematic for decades, they may yet prove essential for understanding the earliest evolution of the Euchelicerata. Once considered part of the monophyletic Xiphosura [10] the synziphosurines have also been considered paraphyletic [11] or even polyphyletic [12]. The polyphyletic reinterpretation fractures the group with some taxa placed as basal to all other euchelicerates while others were scattered around the euchelicerate tree, as sister taxa of chasmataspids, eurypterids and arachnids. In this framework the iconic synziphosurines *Dibasterium durgae* [13] and *Offacolus kingi* [14, 15] have been considered to be stem euchelicerates, possessing defining characters of the modern euchelicerates such as gills opercula and chelicerae. The presence of biramous appendages in the prosoma of *Dibasterium* and *Offacolus* led to the establishment of the clade Prosomapoda, which is the clade of euchelicerates defined by uniramous prosomal appendages [12]. The discovery and description of *Setapedites abundantis* [8] from the Early Ordovician Fezouata Biota [16, 17] united *Dibasterium*, *Offacolus* and *Setapedites* itself into the family Offacolidae [8, 15]. *Habelia optata* [18], previously proposed as the closest Cambrian relative to the euchelicerates [19], was confirmed in its position, on the basis of characters that the Cambrian taxon shares with the expanded Offacolidae (e.g. stenopodous prosomal exopods, uniramous first pair of appendages, bipartite opisthosomal tergites) [8, 19]. Despite the resolved position of Offacolidae, questions remain about certain anatomical characters in *Setapedites*, particularly about the nature of its pretelsonic process [8]. A possible comparable structure was described by Simonetta [20] in *H. optata* as a “perianal plate” closely related with the post ventral plate of *Aglaspis spinifer* [21], but Whittington [22] refuted this hypothesis, interpreting this feature as a taphonomic artefact. New material then allowed Aria & Caron [19] to confirm the presence of a rounded plate below the telson head, which they interpreted as an anal pouch. On the basis of the pretelsonic process [8], a potential relationship to the Vicissicaudata [23] has also been hypothesized for *Setapedites*. A better understanding of synziphosurine anatomy is essential to reconstructing the relationships not only within Euchelicerata, but also between euchelicerates and their potential sister groups, habeliidans and Vicissicaudata.

New specimens assigned to the synziphosurine species *Bunaia woodwardi* [24] from the late Silurian Bertie Lagerstätte (Ontario, Canada) are described herein. These specimens reveal previously unknown anatomical details in their appendages and pretelsonic process. *B. woodwardi* is the youngest offacolid described so far, extending the range of the clade up into the late Silurian and increasing the number of taxa in the clade to four. Detailed preservation of the pretelsonic process allows us to better constrain this character in the Offacolidae and its relation to the anal pouch of *H. optata* and the appendicular derivatives of Vicissicaudata. Vicissicaudata and Euchelicerata have been suggested to be closely related and to form the clade Arachnomorpha (*<u>sensu</u>* Størmer [25]), which has been retrieved as monophyletic by several phylogenetic analyses [4, 19, 26, 27]. This study describes a possible homologous origin for the posterior trunk morphology of Vicissicaudata, Euchelicerata and Habeliida, providing support for a phylogenetic relationship among them and therefore supporting the broader Arachnomorpha hypothesis [25], rather than a monophyletic Artiopoda [28] which does not include Euchelicerata and its sister group the habelliidans.

## 2. Material and methods

### 2.1. Geological Setting

Four specimens were examined as part of this study. All the specimens are housed at the Royal Ontario Museum (catalogued as ROMIP53886, ROMIP51447, ROMIP50689, ROMIP50690). They were collected from two quarries in the Williamsville Formation of the Bertie Group (Piridoli), located near Fort Erie, Ontario, Canada. The Bertie Group is composed of limestones, dolomitic limestones and evaporates, and outcrops are present in western New York State (US) and south-eastern Ontario (Canada). The Williamsville is the Bertie Group’s upper formation, and it consists of fine-grained, argillaceous dolomite of different shades of gray, and dolomitic mudstone [29–32]. The age of the formation is based on correlations with the unit below and above, since the biostratigraphic control for this formation is poor [33].

Specimens ROMIP53886 and ROMIP51447 were collected from the Ridgemount Quarry (42°92’96” N 79°00’48” W), while specimens ROMIP50689 and ROMIP50690 were collected from the George C. Campbell Quarry, approximately 1 mile south of Ridgemount Quarry.

These specimens were compared to photographs of the original four specimens of *B. woodwardi* (catalogued as NYSM 9909, NYSM 9910, NYSM 9911, NYSM 9912). Clarke (1919) refers to this material as originating from the “Bertie Waterlime of the Salina Group” from an outcrop at East of Buffalo (New York state, US). However, following works recognized the Bertie and the Salina as different groups [29–32] so we attribute Clark’s original material to the Bertie Group.

### 2.2. Systematic framework

This work follows the systematics of [4, 8, 12, 19, 23, 34] and uses the anatomical terms presented in [8, 12, 13, 15, 19, 23, 34, 35].

### 2.3. Photography and multispectral imaging

The specimens were each examined with a light microscope (Wild Heerbrugg M8) under normal light. Subsequently they were photographed with an SLR camera (Canon EOS 800D equipped with CANON macro lens MP-E 65 mm 1:2.8 1–5×) mounted on a stand and connected to a z-stacking system (STACKSHOT 3X), using different combination of lightning conditions: normal light, polarized light, dry, covered in alcohol. Z-stacks were rendered using the Helicon Focus software. All specimens were analyzed in the Optical laboratory at Lausanne University (Geopolis-3439). The specimens were also examined under multispectral macroimaging using an innovative setup that allows for the collection of reflection and luminescence images in various spectral ranges from the ultraviolet (UV) to the near infrared (NIR) domains [36–38]. The setup is made of: a low-noise 2.58-megapixel back illuminated sCMOS camera (PRIME 95B 25 mm, Photometrics) with high sensitivity from 200 to 1000 nm, fitted with a UV–VIS–IR 60 mm 1:4 Apo macro lens (CoastalOptics) opposite by a filter wheel holding eight interference band-pass (Semrock) to collect images in eight different spectral ranges from 435 to 935 nm. Illumination with wavelengths ranging from 365 up to 700 nm was provided by 16 LED (CoolLED pE-4000), coupled to a liquid light-guide fibre fitted with a fibre-optic ring light-guide [36–38]. This setup consequently allows for collecting images in a total of more than 90 different illumination/detection configurations. The resulting 16-bit grayscale images were combined into false colour RGB overlays in ImageJ, manually adjusting the mininum and maximum values for each image in order to provide the strongest possible enhancement of contrast, and thus reveal new details of the morphology of the specimens examinated [36–38]. The false colour RGB overlays associated with each of the specimens analyzed were produced using the following configurations: (figure 1*c*,*d*) red— illumination 525 nm/detection 571 ± 36 nm (reflection), green—illum. 470 nm/det. 719 ± 30 nm (luminescence), blue—illum. 385 nm/det. 515 ± 15 nm (lum.); (figure 2*b*) red— illumination 580 nm/detection 935 ± 85 nm (lum.), green—illum. 525 nm/det. 571 ± 36 nm (refl.), blue—illum. 470 nm/det. 719 ± 30 nm (lum.); (figure 3*c*) red—illumination 525 nm/detection 571 ± 36 nm (refl.), green—illum. 470 nm/det. 719 ± 30 nm (lum.), blue—illum. 365 nm/det. 571 ± 36 nm (lum.); (figure 3*d*) red—illumination 365 nm/detection 650 ± 30nm (lum.), green—illum. 770 nm/det. 775 ± 23 nm (refl.), blue—illum. 525 nm/det. 571 ± 36 nm (refl.).

**Figure 1.**
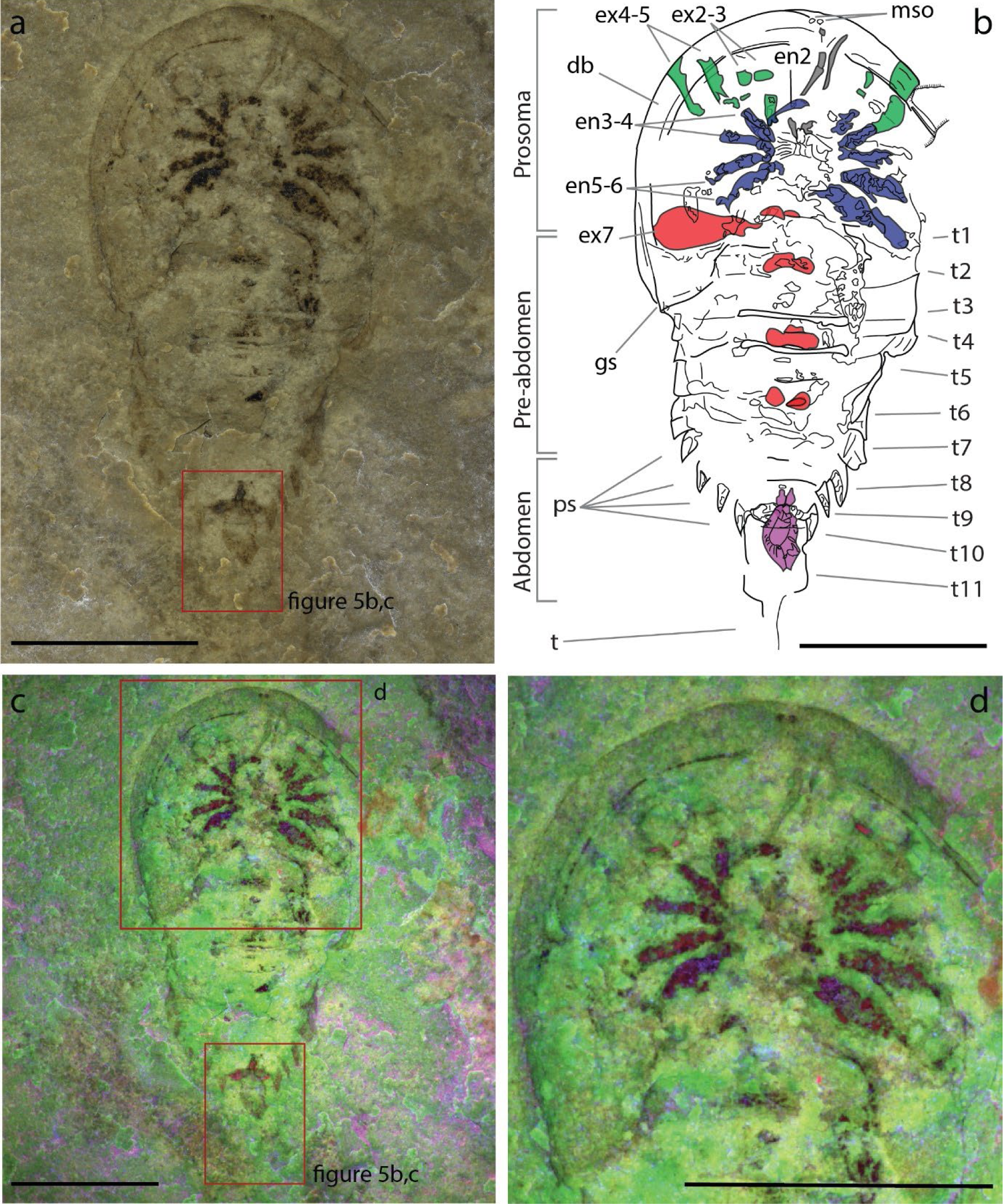
New specimen of *Bunaia woodwardi* ROMIP53886. (*a*) Optical photograph under polarized filter and submerged in alcohol. (*b*) Interpretative line drawing. Highlighted in gray are the chelicerae, in green prosomal exopods, in blue endopods, in red opisthosomal exopods, and in purple the pretelsonic process. (*c*) Multispectral imaging. (*d*) Multispectral imaging close-up from the upper boxed area in (*c*). Abbreviations: en2–5, endopods 2–5; ex2–6, exopods 2–6; mso, median sensory organs; ps, pleural spine; t1–t11, opisthosomal tergites 1–11; t, telson. Scale bars = 1 cm.

**Figure 2.**
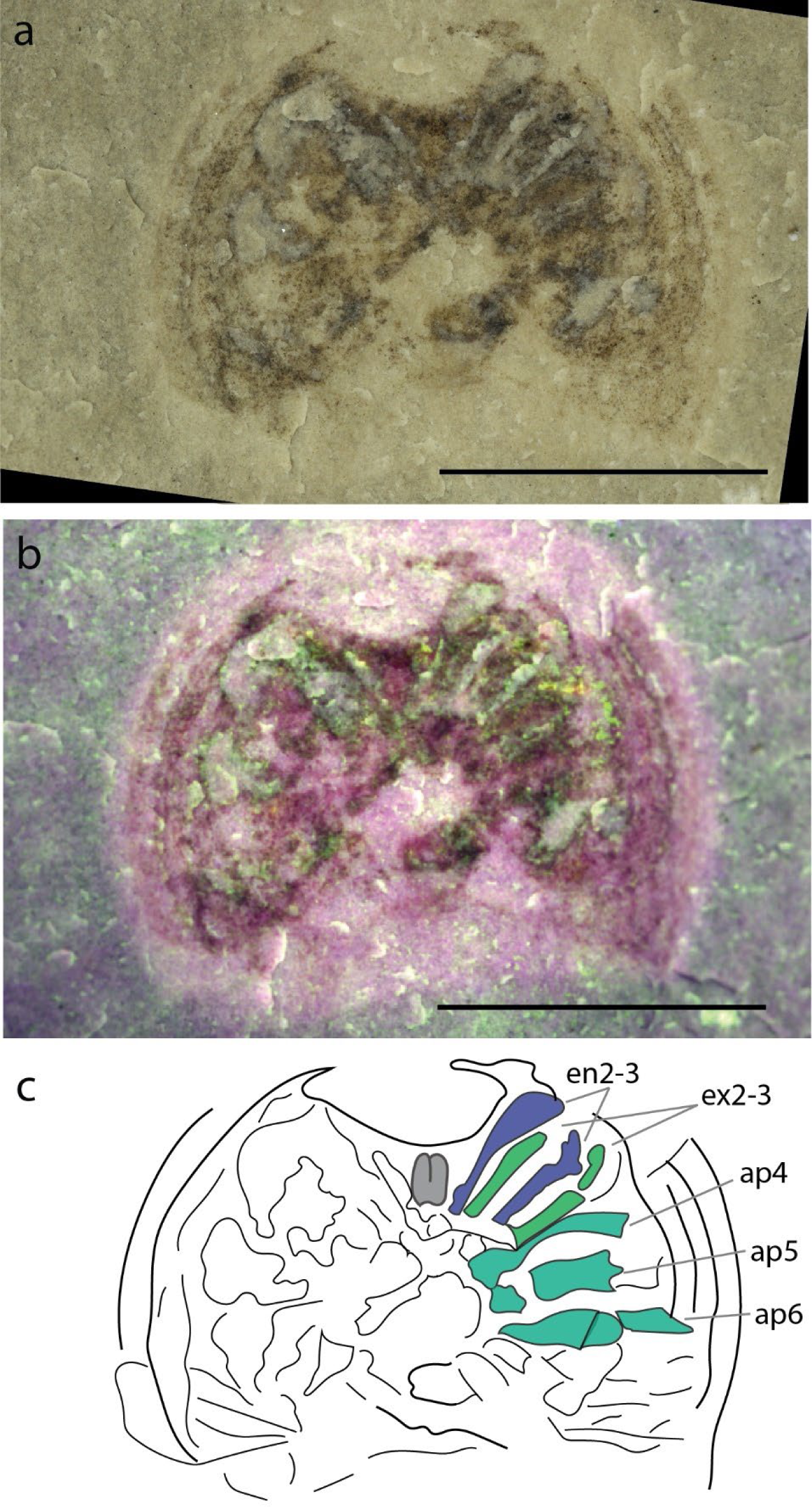
New specimen of *Bunaia woodwardi* ROMIP51447. (*a*) Normal light optical photograph. (*b*) Multispectral imaging. (*c*) Interpretative line drawing. Highlighted in gray are the chelicera, in green prosomal exopods, in blue endopods, in aquamarine unidentified exopods or endopods of appendages 4–6. Abbreviations: ap4–6, appendages 4–6; en 2–3, endopods 2–3; ex2–3, exopods 2–3. Scale bars = 5 mm.

**Figure 3.**
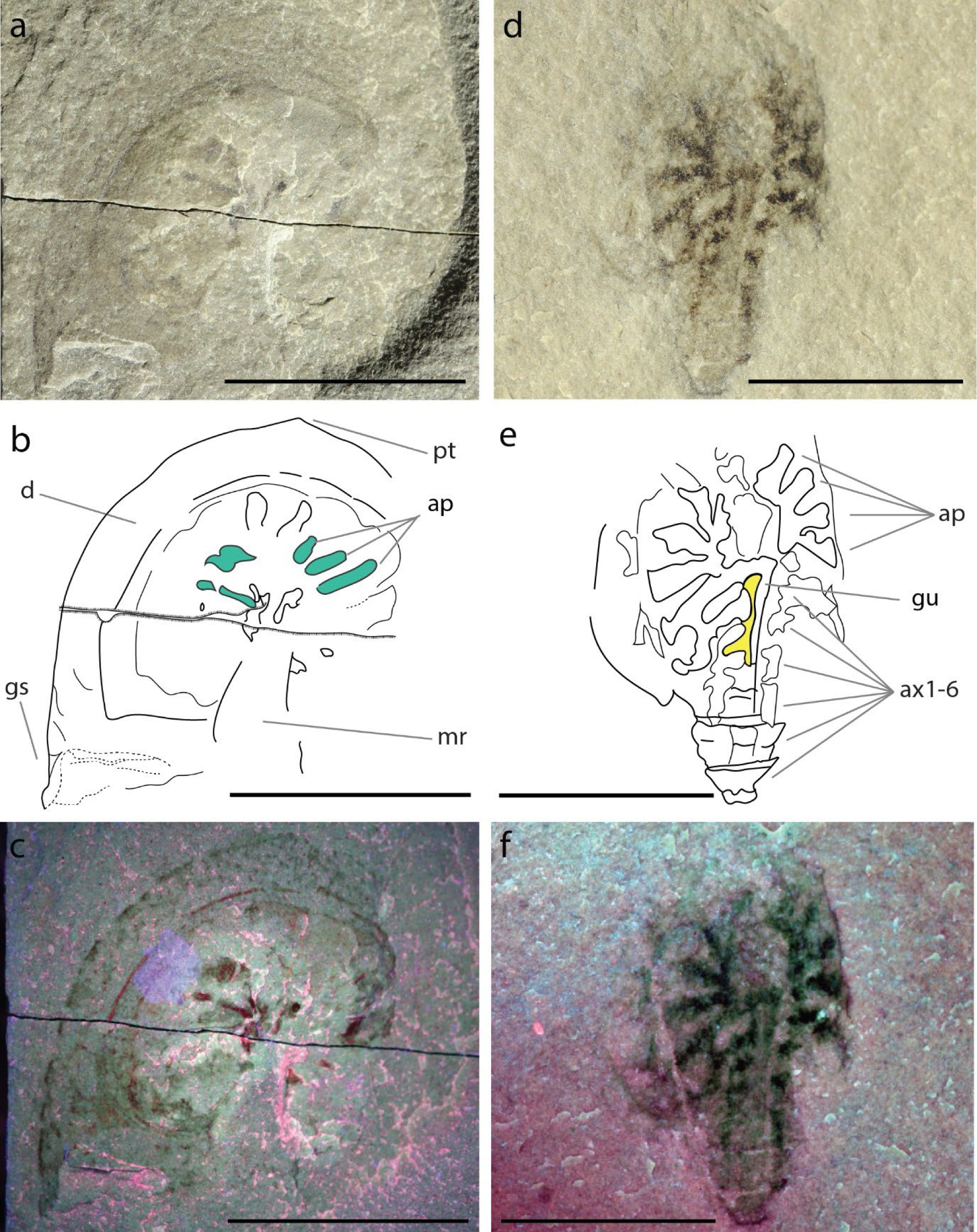
New specimens of *Bunaia woodwardi* ROMIP50689 and ROMIP50690. (*a–c*) Normal light optical photograph (*a*), interpretative line drawing (*b*) and multispectral imaging (*c*) of specimen ROMIP50689. (*d*–*f*) Normal light optical photograph (*d*), interpretative line drawing (*e*) and multispectral imaging (*f*) of specimen ROMIP50690. Highlighted in aquamarine in (*b*) are unidentified exopods or endopods of the appendages, and in yellow, in (*e*) are remains of the digestive system. Abbreviations: ap, appendages; ax1–6, axis 1–6 of the opisthosomal tergites; d, doublure; gs, genal spine; gu, gut; mr, cardiac (median) ridge; tp, apical tip. Scale bars = 1 cm.

### 2.4. Interpretative drawings

Interpretative drawings were made in Adobe Illustrator on the photo, using a Wacom ver. 6.3.38-3 graphic table and a pen tool.

### 2.5. Phylogenetic analyses

Parsimony and Bayesian analyses were performed using the recent standard methodology for this group [8, 12]. *B. woodwardi* was added to the expanded matrix from Lamsdell [12] used in Lustri et al. [8]. Character coding for *B. woodwardi* is available in electronic supplementary material dataset. Parsimony analyses were run on TNT ver. 1.5, using a traditional search with 100 000 random addition sequences followed by branch swapping with 100 000 repetitions. All characters were unordered and both an implied weight (3, 6, 9 and 12K) and an eaqual weight was applied, followed by Bremer support (10000 repetitions). Bayesian analyses were run on mrBayes ver. 3.2.7a using a tree search followed an Mkv + Γ model [39] with four chains sampling during four runs for 10,000,000 Markov chain Monte Carlo generations, a tree sampled every 1,000 generations and burn-in of 20%. The mrBayes script is available as an electronic supplementary material dataset.

### 2.6. Abbreviations

#### 2.6.1. Institutional abbreviations

NIGPAS, Nanjing Institute of Geology and Palaeontology, China; NYSM, New York State Museum, USA; MPM, Milwaukee Public Museum, USA; ROMIP, Royal Ontario Museum Invertebrate Paleontology collections, Canada; UMNH, Natural History Museum of Utah, USA.

#### 2.6.2. Abbreviations used to label figures

ap, appendages; ax, axis; bf, bifurcation; ch, chelicera; mr, median ridge (cardiac ridge); cn, centrum; d, doublure; en2–5, endopods 2–5; ex2–6, exopods 2–6; gs, genal spine; gu, gut; in, insertion; mso, median sensory organs; pd, pleural spine t, telson; t1–t11, opisthosomal tergites 1–11.

## 3. Results

### 3.1. Systematic Paleontology

Arthropoda von Siebold, 1848.

Arachnomorpha Størmer, 1944. Euchelicerata Weygoldt & Paulus, 1979.

Offacolidae Sutton, Briggs, Siveter, Siveter and Orr 2002.

*Bunaia woodwardi* Clarke, 1919.

*. 1919 Bunaia woodwardi* Clarke, pp. 531–532, pl.14, figs. 1–4;

**Figure 4.**
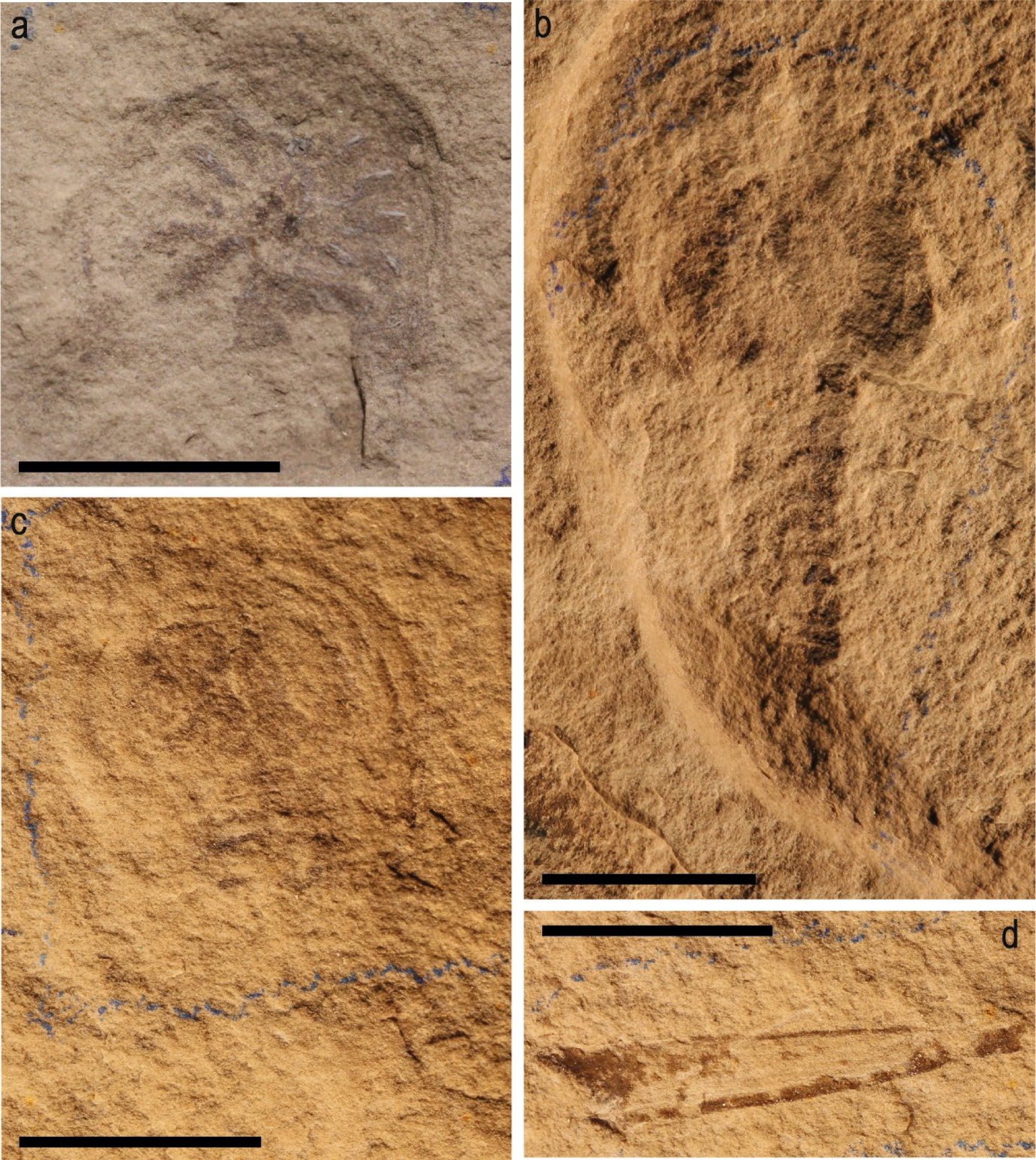
Lectotype and syntypes of *Bunaia woodwardi*. (*a*) Syntype NYSM 9910. (b) Syntype NYSM 9911. (*c*) Lectotype NYSM 9909. (*d*) Syntype NYSM 9912 (all photos credited to Russell Bicknell). Scale bars = 1 cm.

*. 1920 Bunaia woodwardi* Clarke, pp. 129, pl. 1, figs. 1-4;

*. 1925 Bunaia woodwardi* Ruedemann, pp. 79, pl. 24, figs. 1–3;

. 1974 *Bunaia woodwardi* Eldredge and Smith, pp. 24, figs. 8;

*2009 specimen ROMIP53886 figured as *Bunaia woodwardi,* Rudkin and Young, pp. 35, fig. 6 [40].

(figures 1, 2, 3, 4, 5*a*–*c*)

#### Amended diagnosis

Large offacolid species with a total length of up to 32 mm (telson excluded). Prosoma long, about 30% of the total body length (telson excluded). Thick doublure. Eleven opisthosomal somites. Presence of triangular shape pleural spine in opisthosomal somite VII–X. 1^st^ opisthosomal tergite more than four times wider than the 11^th^ tergite, 9^th^ tergite two times wider than the 11^th^.

#### Types

Lectotype, NYSM 9909 (figure 4). Syntypes NYSM 9910, 9911, 9912 (figure 4).

#### Referred material

ROMIP53886 (part; figure 1*a*–*d*); ROMIP51447 (part; figure 2*a*–*c*); ROMIP50689 (part; figure 3*a*–*c*); ROMIP50690 (part; figure *3d*–*f*); NYSM 9909 (figure 4); NYSM 9910 (figure 4); NYSM 9911 (figure 4); NYSM 9912 (figure 4).

### 3.2. Description

#### 3.2.1. ROMIP53886

ROMIP53886 (figure 1) is an almost complete specimen that has been dorso-ventrally flattened during preservation. It possesses an obovate shaped body that is divided into three tagma: an anterior prosoma overlapped by a dorsal headshield, and an unfused opisthosoma not clearly differentiated into a pre-abdomen and an abdomen. Total length of the body (from the tip of the prosoma to the last somite) is 32 mm. The prosoma represents the widest region of the body and its preservation is incomplete on the left side, so the maximum width is estimated to ca. 22 mm.

The headshield is lunate in outline (13 mm long and 22 mm wide). A small genal spine is present on the posterior-left edge (gs in figure 1*b*). A well-developed doublure completely surrounds the prosomal shield ventrally (db in figure 1*b*). No labrum or ophthalmic region can be observed. Two small flat dots medially on the doublure may represent median sensory organs (mso in figure 1*b*).

The prosoma bears six pairs of appendages inserted around the mouth. The first pair of appendages are uniramous, elongate chelicerae, anteroventrally oriented (grey structures in figure 1*b*). However, no evidence for the chelate last podomere is present. Appendages 2 to 5 are biramous and latero-ventrally oriented (en2–6 and ex2–5 in figure 1*b*). Endopods 2 and 3 are bended and allow to recognize the associated exopod 2 and 3(en2–3 and ex2–3 in figure 1*b*). Appendage 6 is possibly uniramous (en6 in figure 1*b*). The endopods are preserved retracted. Details of the anatomy of the podites are not preserved in both external and internal rami.

The prosoma is long, representing about 30% of the total body length (short opisthosoma measures 19 mm, long prosoma measures 13 mm). The opisthosoma consists of eleven somites (somite VII to XVII). It has a trapezoidal shape with the longest side coinciding with the first opisthosomal tergites (somite VII, 3.3 mm wide), and the short side coinciding with the last somite (somite XVII, 1.5 mm wide). A clear boundary between pre-abdomen and post-abdomen is absent. However, a discontinuity in the pleural spines (ps in figure 1*b*) occurs from the seventh somite, underlining the probable border between pre-abdomen and abdomen. The first opisthosomal somite (somite VII) appears reduced, partially or fully overlapped by the prosomal shield and no pleurae are noted. The imprint of the axis is present in somite VIII– XII (opisthosomal tergites 2 to 6), with no sign of antero-posterior bipartite sclerotization noted. No sub axial node is observed. Tergites 1 to 6 possess pleura with a square shape. The width of the tergites decreases slightly from tergites 3 to 10 (from 3.3 mm for tergite 3 to 1.5 mm for tergite 10). Tergites 1 to 10 bear pleural spine.

The first opisthosomal somite (somite VII) bears a pair of appendages (ex6 in figure 1*b*), while somite VIII preserves only the proximal part of them. The first opisthosomal appendages are paddle-like uniramous exopods and insert medially to the somite, in the axial region. Traces of the insertion point of following appendages are also preserved in somites X and XII.

The abdomen is probably composed of four somites (somites XIV to XVII) based on the pleural anatomy, however the absence of preservation of the dorsal anatomy is an obstacle to an unequivocal description. Triangular pleural spines are present on the side of the abdomen tergites (ps in figure 1*b*). Somite XVII is three times longer than the previous somite (somite XVI is 1.5 mm long, somite XVII is 4.5 mm long) and devoid of pleural spines.

A symmetrical process in a pretelsonic position is preserved ventrally to somite XVI–XVII (purple structure in figure 1*b*) (see also figure 5*a*–*c*). This pretelsonic process is longitudinally bipartite and has a bifurcated proximal tip distally (bf in figure 5*c*). It is formed by two branches departing from two points at the margin of somite XVI–XVII (in in figure 5*c*), diverging into an oval shape and meeting again on the distal end, ventrally to the XVII somite. A terminal telson is present, but not completely preserved.

**Figure 5.**
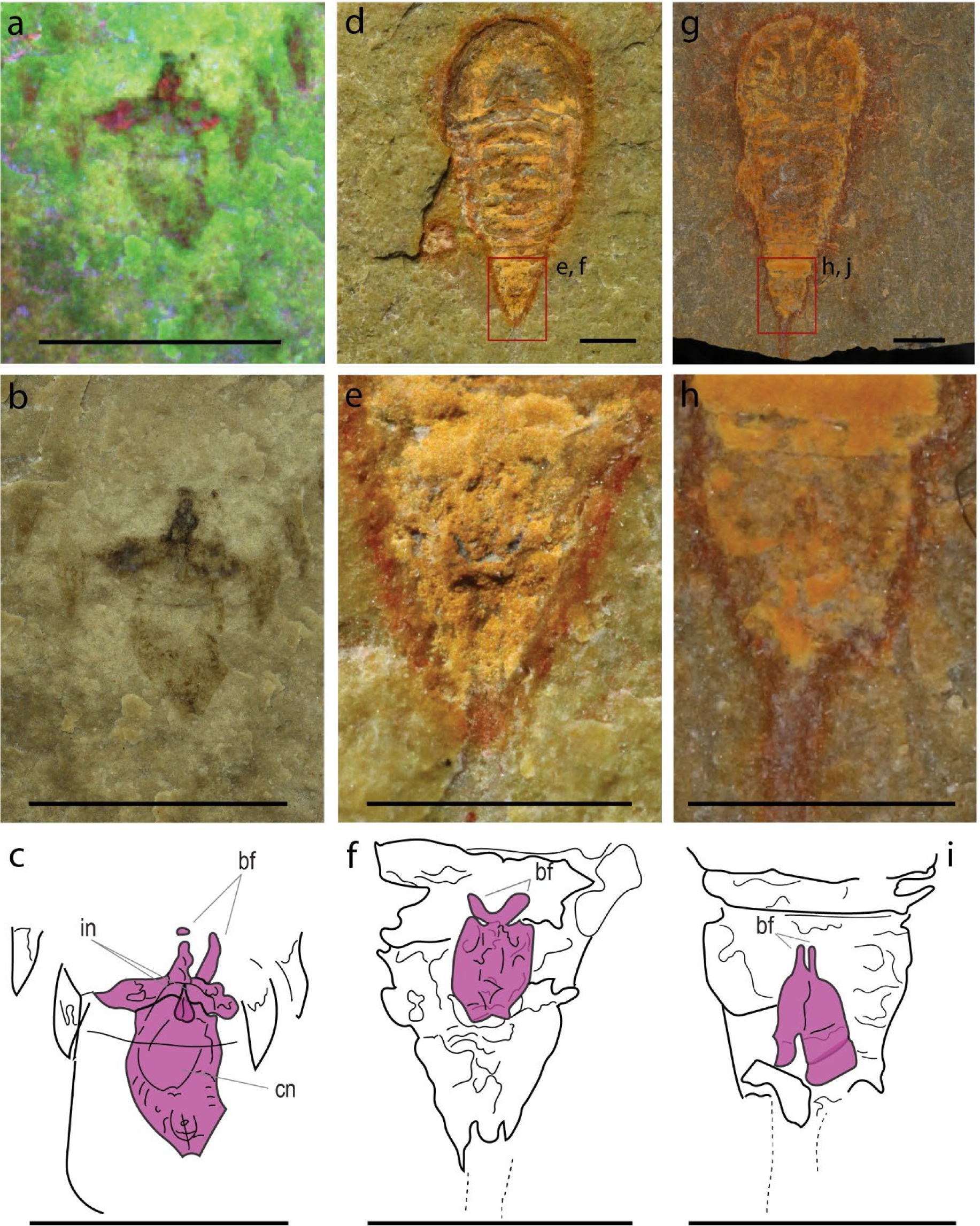
Pretelsonic procesess of *Bunaia woodwardi* and *Setapedites abundantis* (Offacolidae). (*a,b*) Mulispectral imaging (*a*) and optical photography (*b*) close ups of the pretelsonic process of *B. woodwardi* specimen ROMIP53886 from the boxed areas in figure 1c and 1*a*, respectively.. (*c*) Interpretative line drawing of the pretelsonic process of *B. woodwardi* specimen ROMIP53886 based on (*a*,*b*). (*d*–*f*) Habitus view of *S. abundantis* specimen MGL102902 (*d*), and close up (*e*) and interpretative line drawing (*f*) of its pretelsonic process. (*g*–*i*) Habitus view of *. abundantis* specimen YPM IP 517932 (*g*), and close up (*h*) and interpretative line drawing (*i*) of its pretelsonic process. Abbreviations: in, insertion; bf, bifurcation; cn, centrum. The different pretelsonic processes are highlighted in purple. Scale bars = 1 cm (*a*–*c*), 1 mm (*d*–*i*).

#### 3.2.2. ROMIP51447

ROMIP51447 (figure 2*a*–*c*) ventrally preserves only the prosomal region. A well-developed doublure completely surrounds the prosomal shield ventrally. No labrum is observed. The prosoma bears six pairs of appendages that are better preserved on the right side of the specimen. The first pair of uniramous appendages, interpreted as the chelicerae, are retracted. No evidence for the chelate last podomere is present. Appendages 2 to 3 are well preserved, biramous and latero-ventrally oriented. Appendages 4 to 6 are preserved, but a distinction between external and internal rami is not possible. Anatomical details of the podites are not preserved in both exopodite and endopodite.

#### 3.2.3. ROMIP50689

ROMI50689 (figure 3*a*–*c*) also ventrally preserves only the prosomal region. A well-developed doublure completely surrounds the prosomal shield ventrally. An apical tip is present at the anterior edge of the prosoma (pt in figure 3*b*), and a small genal spine is visible on the left side (gs in figure 3*b*). No labrum is observed. The prosomal appendages are preserved (ap in figure 3*b*) but the inner and external rami cannot be distinguished owing to preservation quality. In the middle proximal part of the prosoma the imprint of what could be the median ridge (cardiac ridge) is preserved (mr in figure 3*b*).

#### 3.2.4. ROMIP50690

ROMIP50690 (figure 3*d*–*f*) partially preserves the prosomal and opisthosomal anatomy. It is dorso-ventrally flattened in its preservation. The dorsal prosomal anatomy is completely missing. A general imprint of the appendages is preserved but lacks all anatomical details (ap in figure 3*e*). The opisthosoma preserves six somites but only in the tergites axial anatomy (ax1–6 in figure 3*e*). In the center of the opisthosoma, a straight tubular structure runs along the first three segments. It preserves two lobes departing from the left sides and is interpreted as the digestive system (gu in figure 3*e*). No pleura or opisthosomal appendage anatomy is preserved. Abdomen and telson are not preserved.

### 3.3. Remarks

Two Silurian-aged species of the genus *Bunaia* have been previously described. The first is *Bunaia woodwardi* Clarke [24] from Bertie Group of East Buffalo (New York state, USA) and the second is *Bunaia heintzi* Størmer [41] from Ringerike Sandstone (Norway). The Norwegian material consists of an isolated prosoma, and its assignment to the genus *Bunaia* has previously been questioned. Eldredge [42] pointed out that the differences in prosomal shape, the larger size and the presence of a marked granulation in *B. heintzi* may justify its allocation to a new genus.

Before this formal description of the four specimens from the ROM, *B. woodwardi* was known only from four poorly preserved specimens [24, 42] (figure 4). Specimen ROMIP53886 (figures 1*a*–*c*) was subsequently figured by Rudkin and Young [40] and referred to as *B. woodward*i but not officially described. The specimens are here assigned to species *B. woodwardi* based upon their coeval stratigraphical and geographical occurrence, numerous shared characteristics and the absence of inconsistent characters (see §4.1). We also remark that the genus *Bunaia* itself has been questioned as a possible synonyms of *Pseudoniscus clarkei* [42]. The genus *Pseudoniscus* includes four species from Europe and North America, nominally *Pseudoniscus aculeatus* from the Oesel group in Estonia [43], *Pseudoniscus falcatus* from the Patrick Burn Formation of Scotland [44], *Pseudoniscus roosevelti* from Pittsford shale of New York state (US) [45] and *Pseudoniscus clarkei* from the Bertie Group of New York state (US) [46]. A broad review of the genus *Pseudoniscus* is required in order to assess whether *Bunaia* should be consider a synonym of *Pseudoniscus*.

## 4. DISCUSSION

### 4.1. Attribution of the newly described material to *Bunaia woodwardi*

The exceptional quality of preservation of the new specimens described herein reveals previously unknown anatomical features in great detail, but presents some uncertainty when uniting these specimens (figures 1, 2 and 3) with the original *B. woodwardi* lectotype and syntypes (figure 4) because of the comparably poor preservation of the original type specimens. After the first brief description [24], the four original specimen of *B. woodwardi* (figure 4, NYSM 9909; NYSM 9910; NYSM 9911; NYSM 9912) were re-described in 1974 by Eldredge [42]. This re-description led to the identification of numerous taxonomically significant characters including the presence of a small genal spine, a curved posterior margin of the prosoma, the presence of a cardiac ridge, and the median sensory organs [42]. Most of these anatomical features are also observed in the specimens housed in the ROM. The small genal spine and curved posterior margin of the prosoma are seen in ROMIP53886, the presence of a cardiac ridge (median ridge) is suggested by ROMI50689, and possible median sensory organs have also been identified in ROMIP53886 (figures 1*a*–*c*, 3*a*–*c*). Other characters shared among all the specimens include an expanded doublure (ROMIP51447, ROMIP53886, ROMIP50689, NYSM 9909; respectively figures 1, 2, 3*a*–*c*, 4*c*) and an opisthosomal axis (ROMIP50690, NYSM 9911; respectively figures 3*d*–*f*, 4*b*). These characteristics taken together substantially contribute to uniting these specimens with the species *B. woodwardi*. However, most of these characters are also commonly found in other synziphosurine taxa e.g. *Legrandella lombardii*, *Bunodes lunula, Cyamocephalus loganensis* and *Weinbergina opitzi* [42, 47–49] and as such are of weak broader taxonomical relevance. Other factors support placing the ROM specimens within *B. woodwardia*. Firstly, all the specimens originated from the Bertie Formation, even if from two different localities (Ridgemount Quarry in Ontario and East Buffalo in New York State). The specimens are not disparate in time and space, only in preservation quality. Secondly, none of the original paratype specimens preserve traces of appendages, the pretelsonic process, or the abdominal pleural spine that contradict what can be identified in the better preserved and newly described material herein [24, 42] (figures 1–4). Therefore, it is necessary to balance the shared anatomy, the coeval stratigraphy, and the geographical proximity of the fossil sites against the lack of more specific diagnostic characters in the original type material. *P. clarkei* [46], also from the Bertie Group, has been considered for the assignation of the newly described material, however, a revision of the entire genus is required before it is possible to compare it with the material here ascribed as *B. woodwardi*, but their close affinities are noted. In balance, it seems reasonable to assign the new material and their unique features to the existing species *B. woodwardi* while providing an emended diagnosis for the species based on the new better-preserved specimens.

### 4.2. *B. woodwardi* is an offacolid

All phylogenetic analyses conducted under parsimony (using both equal weight, see electronic supplementary material, figure S1, and implied weight with an array of different K values, see electronic supplementary material, figures S2-4 for K=3, K=6, K=9 and figure 6a for K=12) and Bayesian methods (figure 6b) retrieved *B. woodwardi* as a member of the family Offacolidae. Prosomal appendages are crucial characters to untangling the sequence of early euchelicerate evolution: the clade Prosomapoda is defined as euchelicerates bearing uniramous prosomal appendages [12], whereas the sister clade of prosomapods, the Offacolidae [15], is defined by the presence of biramous prosomal appendages [12]. These two monophyletic groupings, the Prosomapoda and Offacolidae, collectively comprise the monophyletic group of Euchelicerata [8] (figure 6).

**Figure 6.**
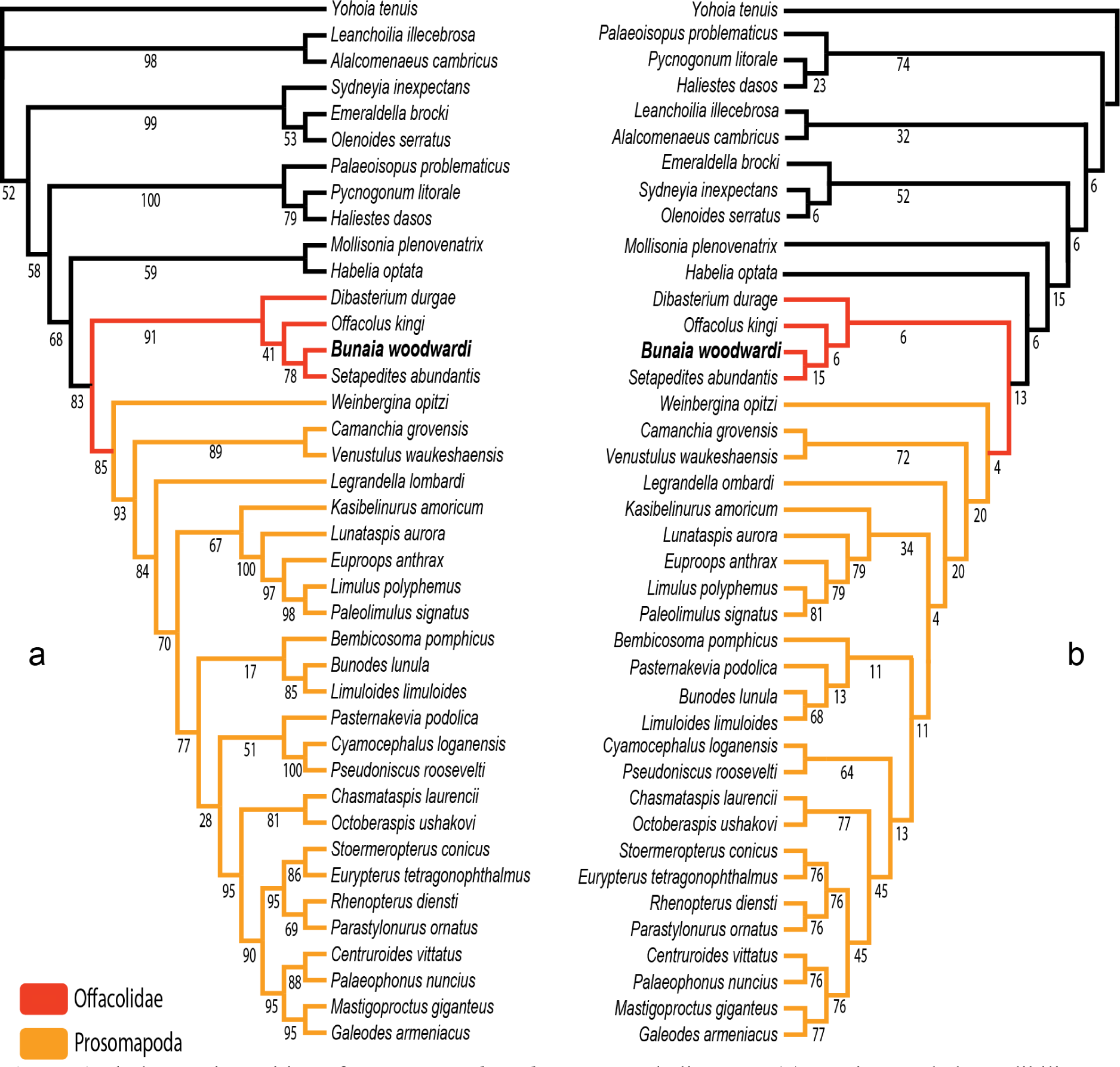
Phylogenetic position of *Bunaia woodwardi* among euchelicerates. (*a*) Maximum clade credibility tree of Bayesian analysis of euchelicerates. Numbers next to nodes are posterior probabilities when <100. (*b*) Implied weights K= 12 maximum parsimony tree of euchelicerates. Numbers next to nodes are relative Bremer support. Both trees are based on a matrix of 40 taxa and 114 characters modified Lamsdell [12] and Lustri *et al.* [8].

The new specimens attributed to *B. woodwardi* (figures 1–3) described herein possess their own suite of diagnostic characters (size, thick doublure, morphology of the abdomen) and also show compelling anatomical affinity to the Offacolidae [8, 13–15]. *B*. *woodwardi* shares three major distinctive characters with other Offacolidae. Firstly, the biramous prosomal appendages with stenopodous exopods [8], an apomorphy that the Offacolidae share with the Cambrian habellidians [19, 26]. Secondly, the elongate uniramous first pair of appendages (chelicerae). Thirdly, the uniramous and paddle-like pair of post-cheliceral appendage 6, apomorphic of the Offacolidae [8]. In addition to these, *B*. *woodwardi* also shares with *Setapedites*, the Fezouata Offacolidae, a pretelsonic process [8], making this structures another possible apomorphy of the Offacolidae. Other distinctive plesiomorphic characters include the presence of a doublure, a moderate to highly developed axis, the number of appendages in the prosoma, and the reduced first tergites, all of which are found preserved in the specimens of *B. woodwardi* (figures 1–3) and suggest this species is a member of Offacolidae. The detailed anatomy of the stenopodus exopods found in *B. woodwardi* are obscure (figure 1). Further fossil specimens are needed to help clarify the detailed anatomy of this key character for the Offacolidae. As stem-euchelicerates, the chelicerae are considered plesiomorphic for Offacolidae. However, the characteristic elongated chelicerae visible in figure 1 are likely a derived character of offacolids. The elongate morphology and the high number of segments of these chelicerae, as shown in *Dibasterium*, make them morphologically similar to the homologous antenna [13, 50, 51]. The morphology of the first pair of appendages in the euchelicerates is similar to that found in their possible close relatives, such as the great appendage of the megacheirans [52], the uniramous short appendages of habeliids [19] and the chelifores of pycnogonids [53]. However as the interrelationships between these major groups remain unclear, it is not yet possible to trace the evolutionary path that led from antenna-like deuterocerebral appendages [50] to the chelate and elongated chelicerae of offacolids. The relationship of offacolids with habeliids [8, 19] is sustained in this work and this suggests that the evolution of the chelicerae occurred earlier in the evolution of Euchelicerata, followed by a diversification early in the group history, with derived forms found in the Prosomapoda and the Offacolidae.

An expanded cephalic doublure is present in *B. woodwardi* (figures 1, 2, 3*a*–*c*) as well as in *Setapedites,* which are respectively the oldest and youngest representatives of Offacolidae. *Dibasterium* and *Offacolus* do not possess a cephalic doublure [13–15]. The cephalic doublure is considered to be of limited phylogenetic importance as it is plesiomorphic and found in numerus Cambrian arthropods such as *Sidneyia inexpectans* [54–56], *Emeraldella brocki* Walcott, 1912 [57, 58], aglaspidids [59] and all trilobites [60]. Among Palaeozoic euchelicerates, the doublure is commonly present among Xiphosura [61] and also in synziphosurines [42, 48]. The presence of a doublure in the Offacolidae adds value to the interpretation of it as a plesiomorphic condition for the whole of the Euchelicerata, and its loss is derived in *Offacolus* and *Dibasterium,* although a homoplastic evolution of this character in *Setapedites* and *B. woodwardi* after its loss at the root of Offacolidae cannot be excluded.

Finally, when comparing the pretelsonic process of *Setapedites* and *B. woodwardi* (figures 1*a*–*c*, 5*a*–*c*) it can be seen that both of these processes are found in the pretelsonic somite and with an attachment point on the border of the previous somite. The shapes are similar, with a horizontal bilateral symmetry expressed by a midline in the middle of the process and a characteristic bifurcation proximal to the insertion point. The newly described pretelsonic process in *B. woodwardi* not only solidifies the interpretation of the same character in the related *Setapedites*, but also suggests that the pretelsonic process might be an apomorphic character of the family Offacolidae. The published descriptions of *Offacolus* and *Dibasterium* have not reported the presence of any particular posterior trunk morphology, however the thin sections-based 3D computerized reconstruction of *Offacolus* specimen OUM C.29557 ([15], fig. 1*a*,*g*) shows a rounded process on the ventral surface below the pretelsonic tergite and preceding the telson. A re-evaluation of *Offacolus* and *Dibasterium* is needed to establish the nature of this structure and its relation to the pretelsonic process of other offacolids. Otherwise, a secondary loss of this character could also explain its absence in both *Offacolus* and *Dibasterium*.

### 4.3. Nature of the different posterior trunk morphologies and their implication for the Arachnomorpha

#### 4.3.1. Posterior trunk morphology

Arachnomorpha (*<u>sensu</u>* Størmer [25]) is the monophyletic group of arthropods composed of Artiopoda (*<u>sensu</u>* Hou [28]) and Chelicerata. This group has been recovered in different phylogenetic analyses with different approaches [4, 19, 27]. Among the Artiopoda, the two main divisions are the Trilobita and the Vicissicaudata [23, 34, 35, 62]. However, the affinity of the Artiopoda with the Chelicerata is not always recovered in phylogenetic analyses [23, 63]. In this phylogenetic framework, Vicissicaudata is a monophyletic group of taxa with a variety of differentiated posterior trunk morphologies, united by the presence of appendicular derivatives sharing a homologous origin [23, 34, 62]. Even though this character can be secondarily lost, as observed in *Carimersa neptuni* [64], the appendicular derivatives are the most distinctive features uniting the group [34]. Vicissicaudata includes the genera *Emeraldella* [58, 65] and *Sidneyia* [56], which possess a uropod and caudal flap (figure 7*a*–*c*); the Aglaspidida [18], which possess a postventral plate or a furcal rami (figure 8); and the Cheloniellida [66], which possess furcae [67] (The terminology used for the posterior trunk morphology in this paper and in the literature is summarized in Table 1).

**Figure 7.**
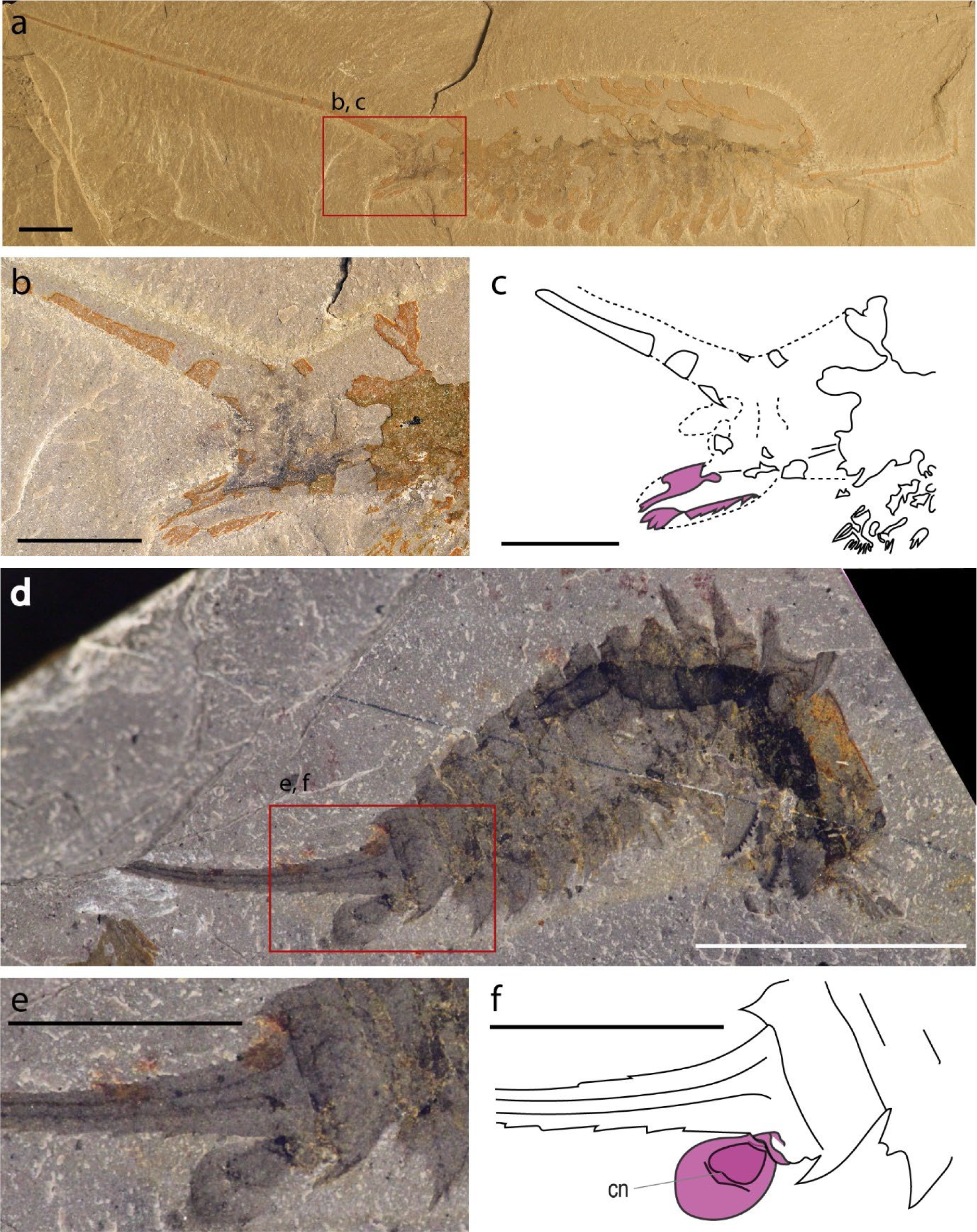
Appendicular derivatives of *Emeraldella brutoni* (Vicissicaudata) and the anal pouch of *Habelia optata*. (*a*–*c*) Habitus view of *E. brutoni* specimen UMNH.IP.6162 (*a*), and close up (*b*) and interpretative line drawing (*c*) of its pretelsonic appendages. (*d*–*f*) Habitus view of *H. optata* specimen ROMIP 64369P (*d*), and close up (*e*) and interpretative line drawing (*f*) of its anal pouch. Abbreviations: cn, centrum. Appendicular derivatives are highlighted in purple. Photography credits: Javier Ortega-Hernández (*a*,*b*), Cédric Aria (*d*,*e*). Scale bars = 5 mm.

**Figure 8.**
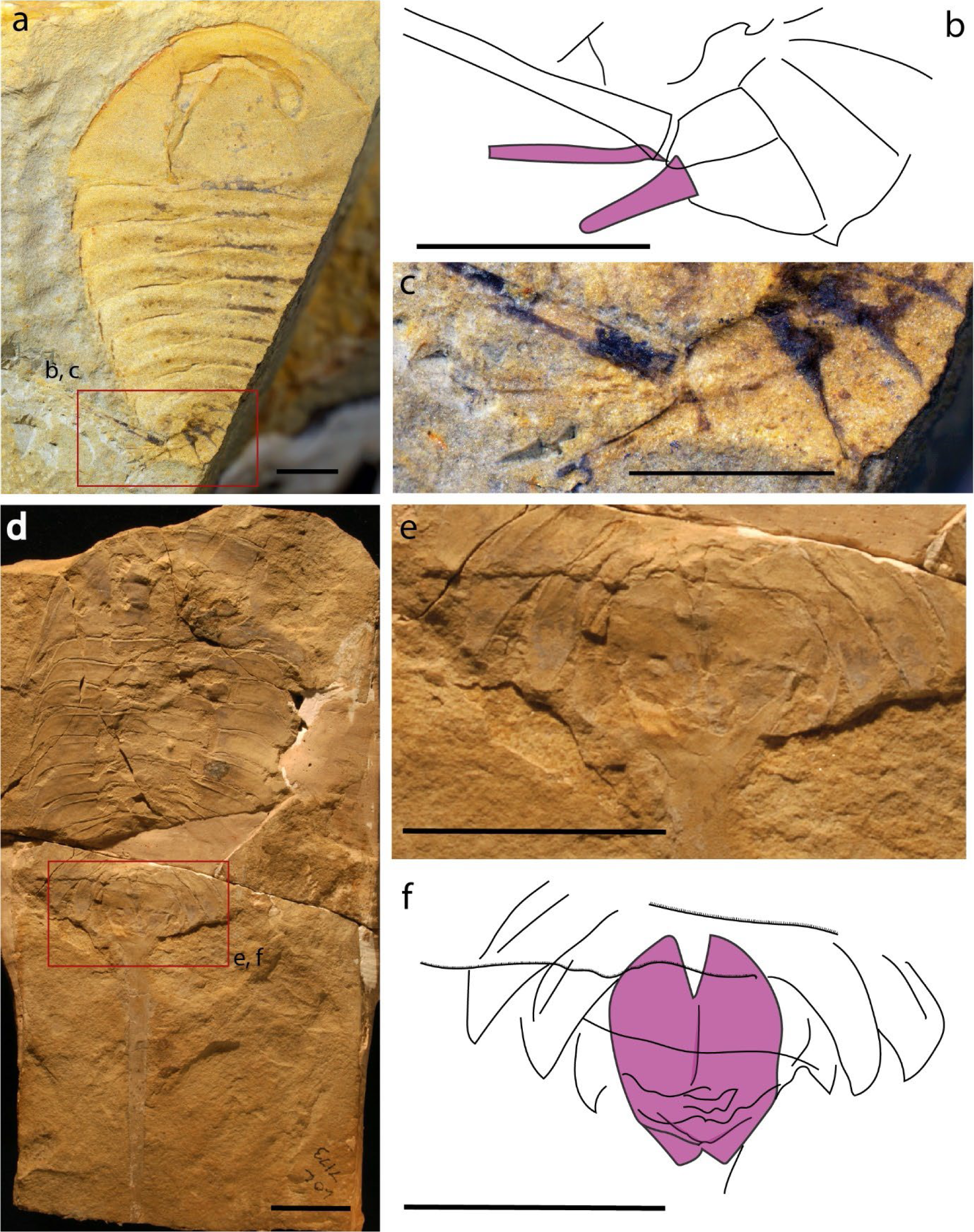
Appendicular derivatives of Aglaspidida. (*a*–*c*) Habitus view of *Glypharthrus tridentatus* specimen NIGPAS165042a (*a*), and close up (*b*) and interpretative line drawing (*c*) of its pretelsonic appendages. (*d*–*f*) Habitus view of *Aglaspis spinifer* specimen MPM11155 (*d*), and close up (*e*) and interpretative line drawing (*f*) of its pretelsonic appendages (postventral plate). The different appendicular derivatives are highlighted in purple. Photography credits: Xuejian Zhu (*a*,*b*), Javier Ortega-Hernández (*d*,*e*). Scale bare = 1 mm (*a*–*c*), 2 mm (*d*–*f*).

**Table 1.**
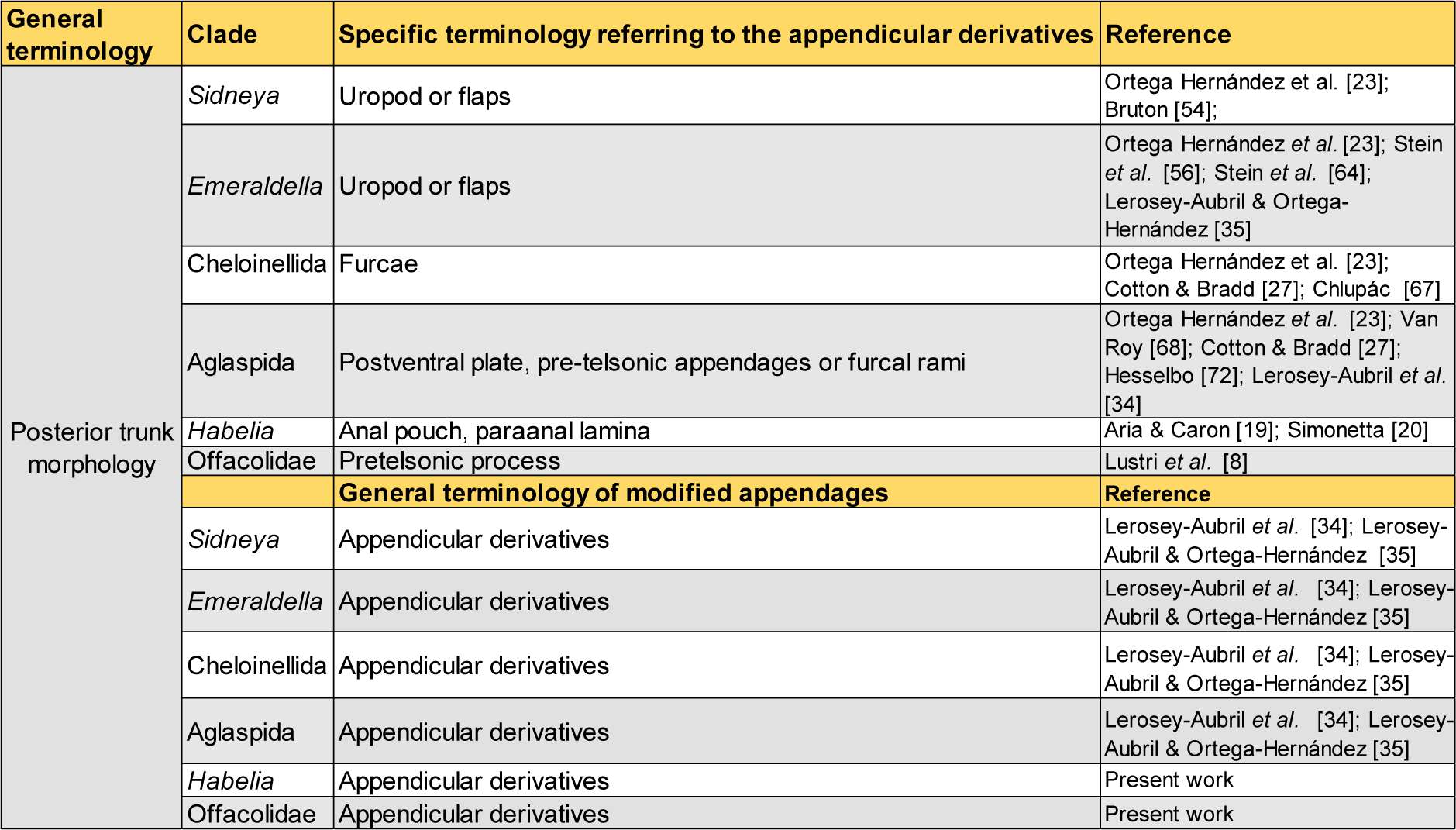
The terminology used to refer to the different posterior trunk morphologies and their appendicular derivatives.

To better understand the Vicissicaudata, several studies have tried to establish the homology of the different posterior trunk morphologies, focusing on the appendicular derivatives and last two somites involved [23, 27, 34], using mainly a positional criterion [68] and acknowledging the fact that all these structures are paired [27, 69]. Indeed, those differentiated appendicular derivatives seem to be always located on the last somite of the abdomen, even though in the evolution of the postventral plate and the furcal rami of the Aglaspidida an alternative hypothesis has been proposed, with the postventral plate possibly originating from the second pretelson somite (tergite 11) and the furcal rami from the pretelsonic somite (tergite 12), meaning they are not homologous [34]. The discovery of *Setapedites*[8] from the lower Ordovician Fezouata Lagerstätte [16, 17], however, suggested that the family Offacolidae may share a homologous posterior trunk morphology with the Vicissicaudata, a hypothesis that if confirmed would add strong support to the Arachnomorpha as a monophyletic group. In particular, the pretelsonic process of *Setapedites* (and now *B. woodwardi*) would be homologous to the appendicular derivatives characteristic of the Vicissicaudata posterior trunk morphology (Supplemental discussions in [8]). The anal pouch of *H. optata* (figure 7*d*–*f*) has also been considered as a possible homologue to the pretelsonic process of *Setapedites* (Supplemental discussions in [8]), because Offacolidae and Habeliida have been recovered as closely related groups in phylogenetic analyses and because of the numerous characters shared between the groups [8, 19, 26].

The following three main interpretations have been suggested (Supplemental discussions in [8]). First, that the pretelsonic process of offacolids is an anal pouch like that in *H. optata*. Second, that the pretelsonic process of offacolids is homologous to the appendicular derivatives characteristic of the Vicissicaudata. And third, if both the previous interpretations are correct, all the supposed appendicular derivatives (*H. optata* anal pouch, Vicissicaudata appendicular derivatives and Offacolidae pretelsonic process) are homologous and reflect the apomorphic state of the inner branches of Arachnomorpha (figure 9). This can be resolved by examining the nature of the posterior trunk morphologies of the groups involved.

**Figure 9.**
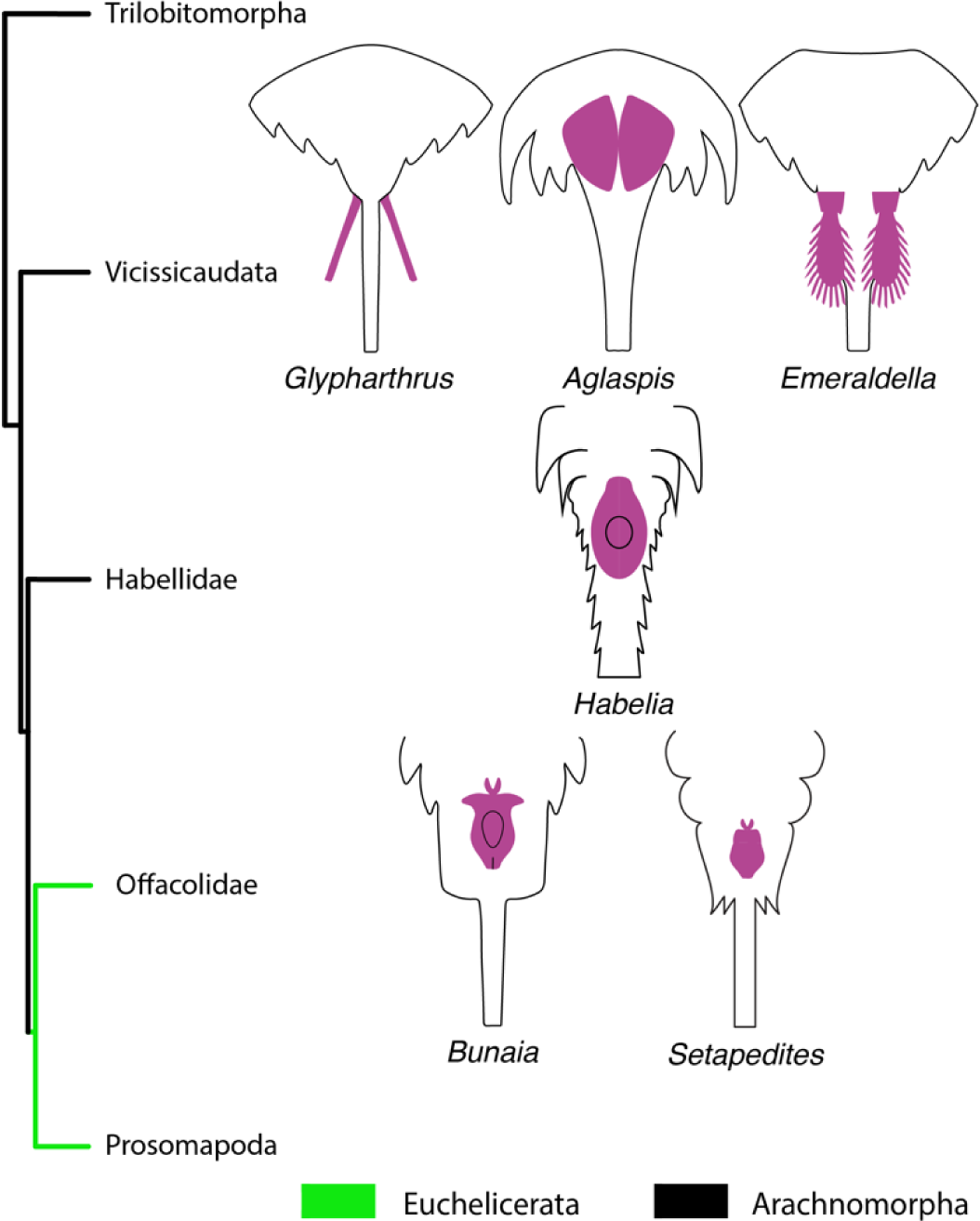
Schematic reconstructions of different posterior trunk morphologies of *Habelia*, Vicissicaudata and Offacolidae plotted onto a hypothetical phylogenetic relationship of Arachnomorpha. Phylogenetic tree based upon the present study results and [4, 8, 12, 34]. Schematic reconstructions drawn from [8, 19, 23]. The different appendicular derivatives are highlighted in purple.

#### 4.3.2. Pretelsonic process and its homology with Habeliida and Vicissicaudata posterior trunk morphologies

The new specimens of the offacolid *B. woodwardi* are well preserved in the posterior trunk morphology, which allows comparison of this anatomical character among the Offacolidae, the Habeliida and the Vicissicaudata (mainly Aglaspidida and the genus *Emeraldella*) in greater detail than has been possible previously (figures 6–9). The posterior trunk morphology of the Cheloniellida is herein excluded from the comparison because although it possesses a pair of furcae, these are located on the dorsal side of the posterior trunk whereas all the other members shows the modified appendages ventrally located [23, 34, 35, 54, 59], suggesting the dorsal location is a derived character of Cheloniellida [70].

Three main criteria are used to establish homology among different characters: (1) shared development, (2) traceable ancestry for derived characters, and (3) association criterion [71]. A developmental pattern regarding the posterior trunk morphology is missing in the whole clade, so the first criterion is excluded from our comparison. Similarly, the uncertainty of the phylogenetic relationship among all the clades of interest makes it difficult to discuss the second criterion. This leaves only the third criterion to consider, the position and symmetry of the different posterior trunk morphologies, to establish homology. However, general morphological considerations such as the degree of sclerotization and overall anatomy are also discussed.

First, the position is considered. The pretelsonic process in *B. woodwardi* is clearly shown in specimen ROMIP53886 (figures 1, 5*a*–*c*). In *H. optata*, the position of the anal pouch is not completely clear. In figure 7 *d*–*f* the anal pouch appears to be inserted in the telson head, while specimen ROMIP 64368 shows a more proximal insertion [19]. In the Vicissicaudata the appendicular derivatives have been associated with the pretelsonic somite, or to the previous somite if the postventral plate is not homologous to all the other appendicular derivatives [34]. In the Offacolidae, the two examples of pretelsonic processes in *Setapedites* and *B. woodwardi* share the same position on the pretelsonic somite (figures 1, 5). With the exception of the Ordovician aglaspidids, which may have had an independent evolution of the postventral plate, the position of all the other appendicular derivatives, including *H. optata* and offacolids, support a shared evolution.

Secondly the bilaterally symmetrical nature of the pretelsonic process is also clear in the Offacolidae and this helps to resolve the position of the homologues with respect to the rest of the anatomy. The double insertion of the *B. woodwardi* pretelsonic process (figure 5*a*–*c*) likely represents its vestigial paired appendicular nature. The modified appendicular derivatives of Aglaspidida, *S. inexpectans* and the genus *Emeraldella* also have a bilateral symmetry. This strongly supports the hypothesis of the pretelsonic process having derived from a pair of appendages inserted in the last post-abdominal segment and being subsequently fused into one single flap. This model is compatible with the evolution of the appendicular derivatives especially in the Aglaspidida, which similarly shows the two plates attached by a less sclerotized membrane [72]. The homology between the pretelsonic process of Offacolidae and the anal pouch of *Habelia* is more difficult to establish in the context of bilaterally symmetry. While Simonetta [20] interpreted the posterior trunk morphology of *H. optata* as a “perianal plate” closely related with the post ventral plate of the Aglaspidida, more recently this process was reinterpreted as an anal pouch that is possibly related to sexual dimorphism [19]. However, in the anal pouch model no possible homologous features in related taxa were identified. Further study on the origins and affinities of the anal pouch in *H. optata* is required, however it currently cannot be excluded to have had originated from appendages that subsequently fused into a single unit.

A general consideration of the degree of sclerotization in the pretelsonic processes is also useful in establishing a homologous origin. Multispectral imaging data show a comparable chemistry between *B. woodwardi* appendages and its pretelsonic process (figure 1*c*), suggesting light sclerotization of the pretelsonic process, as seen in the prosomal appendages. In *Setapedites,* the rare preservation of the pretelsonic process, seen in only five specimens out of many hundreds, suggests a weakly sclerotized or even membranous nature for this structure [8]. This is probably also the case for the habeliids [19], and some of the Vicissicaudata, such as *Emeraldella brutoni*, which has a caudal flap of the same shape and with the same preservation as the exopods [35]. However, Aglaspida present a high degree of sclerotization in the post ventral plate. This suggests that light sclerotization could be plesiomorphic for all the appendicular derivatives, with the high degree of sclerotization in the posterior ventral plate of aglaspidids being a derived state amongst all the appendicular derivatives.

Finally, the flap-like anatomy of the pretelsonic process in *B. woodwardi* can be recognized in the anal pouch of *Habelia*, in that both share a similar rounded or sub-oval shape and a characteristic rounded centrum (see figures 5, 7*d*–*f*, 9 for comparison). Other appendicular derivatives do not show a similar sub-oval shape and a rounded centrum. The close affinities of *H. optata* and offacolids [8, 19] explain this unique combination of characters in their appendicular derivatives as apomorphic traits for *H. optata* and Offacolidae.

## 5. CONCLUSION

New anatomical data described in *B. woodwardi* allow for the placement of this taxon within the family Offacolidae and solidifies the status of this family as a monophyletic group of stem-euchelicerates. The diversity of Offacolidae is now known to be higher, with greater disparity of the abdomen relative size and morphology, and the temporal range of the clade has been extended (by almost 15 million years) to the end of the Silurian. Our reevaluation of synziphosurine taxa using new methodologies (multispectral imaging) allows for a more detailed description of the ventral anatomy. Considering the position and symmetry as well as the general morphology, the appendicular derivatives in the Vicissicaudata and *H. optata* are likely homologous to the pretelsonic process in the Offacolidae, meaning that these taxa share a homologous posterior trunk morphology. However, the lack of a fully resolved phylogenetic framework remains problematic when attempting to establish homology. Further study on the anal pouch of *H. optata* and new phylogenetic studies accounting for these characters would confirm or deny whether posterior trunk morphologies in the Vicissicaudata and Offacolidae share an ancient appendicular origin. A possible phylogenetic framework with graphical schematic reconstructing of all the discussed appendicular derivatives is presented in figure 9. The anatomical analyses of posterior trunk morphologies herein support the Arachnomorpha as a monophyletic group that includes Euchelicerata, Vicissicaudata and Habeliida and help to resolve the highly enigmatic phylogeny at the base of the euchelicerates. This has resulted in much better resolution in the Euchelicerate stem lineage and the sequence of character acquisition during early euchelicerate evolution.

## Data accessibility

XXXXX

## Authors’ contribution

L.L. photographed and drew the studied fossils, performed the phylogenetic analyses, prepared the figures and wrote the manuscript under the supervision and editing of J.B.A., P.G. and A.C.D. P.G. collected and processed the multispectral imaging data. All authors read and approved the final manuscript.

## Competing interests

The authors have no competing interests.

## Funding

L.L. and this research were funded by the Swiss National Science Foundation, grant number 205321_179084 entitled “Arthropod Evolution during the Ordovician Radiation: Insights from the Fezouata Biota” awarded to A.C.D.

## Acknowledgements

We thank Maryam Akrami and Jean-Bernard Caron for shipment of the studied material. We are grateful to Javier Ortega-Hernández, Cedric Aria, Xuejian Zhu and Russell Bicknell for providing photos. Rudy Lerosey-Aubril shared comments and helped with photo acquisition. Peter Van Roy and Javier Ortega-Hernández are also thanked for their discussion of this work.

---

**Supplementary Figure 1.**
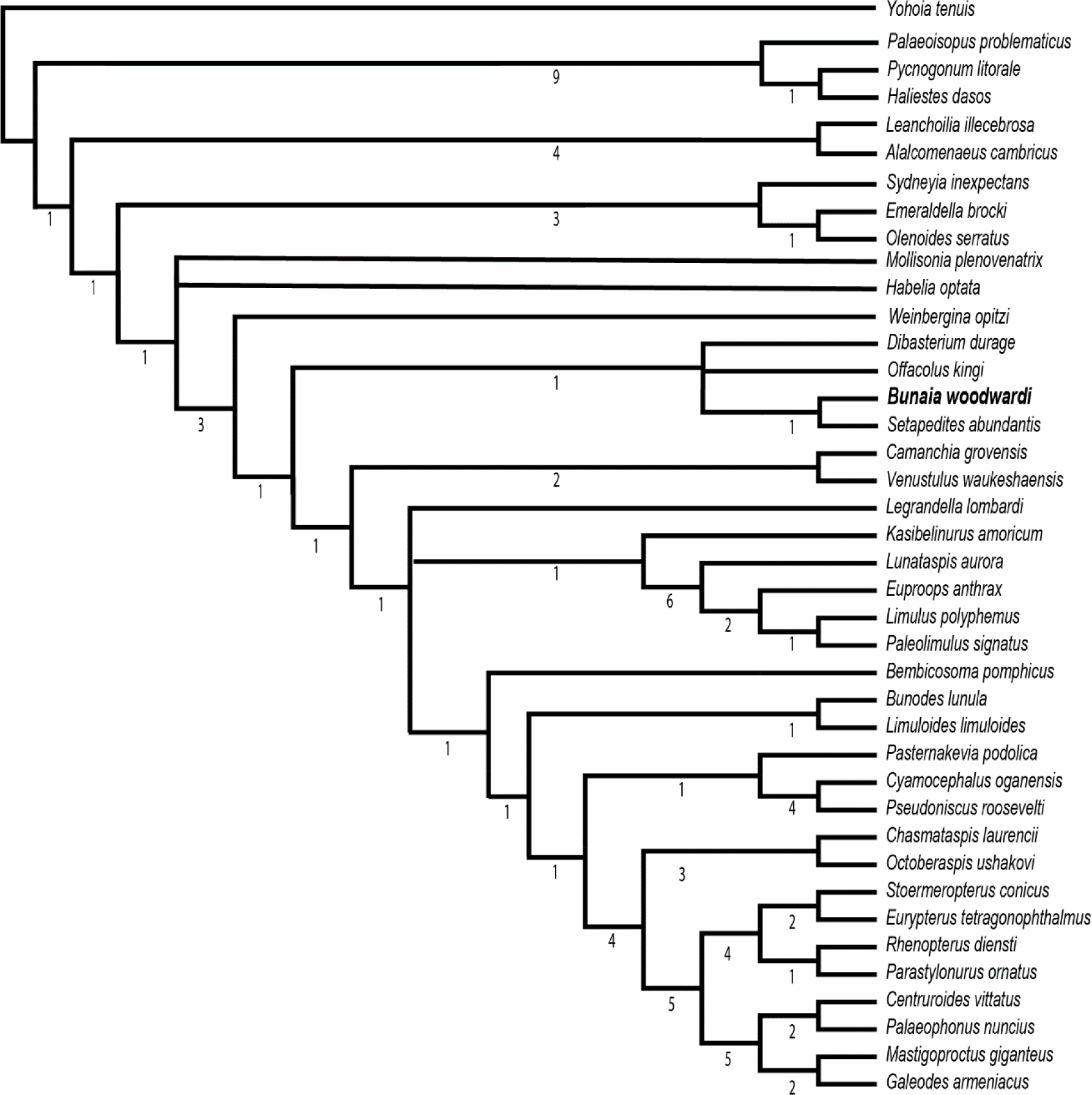
Phylogenetic position of *Bunaia woodwardi* among euchelicerates. Parsimony phylogenetic analysis performed using random addition sequences followed by branch swapping with 100 000 repetitions, all characters unordered and of equal weight. Bremer support is shown with no brackets at the nodes. Matrix modified from Lamsdell [12] and Lustri *et al.* [8].

**Supplementary Figure 2.**
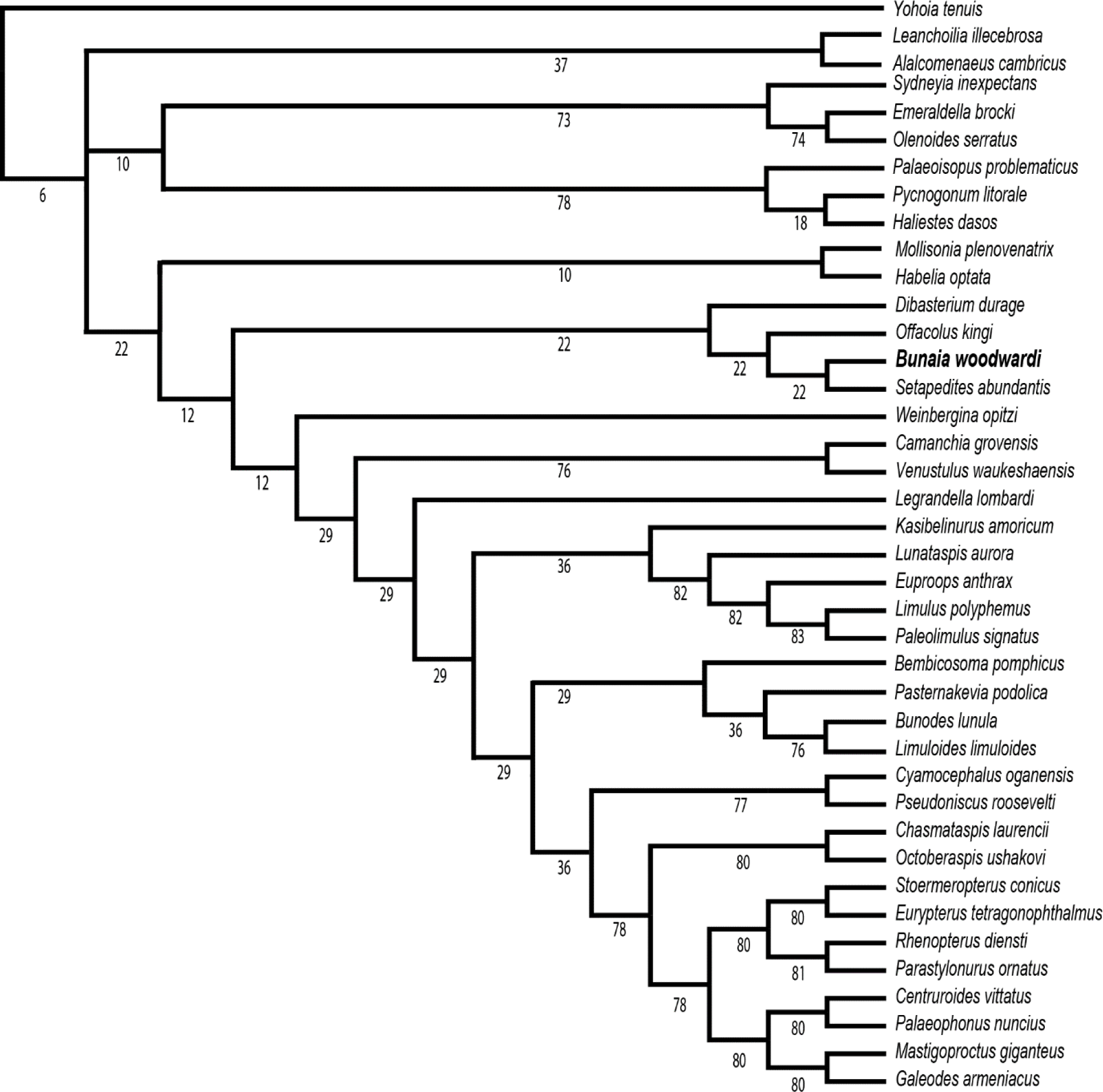
Phylogenetic position of *Bunaia woodwardi.* among euchelicerates. Parsimony phylogenetic analysis performed using random addition sequences followed by branch swapping with 100 000 repetitions, implied weight K= 3. Relative bremer support is shown with no brackets at the nodes. Matrix modified from Lamsdell [12] and Lustri *et al.* [8].

**Supplementary Figure 3.**
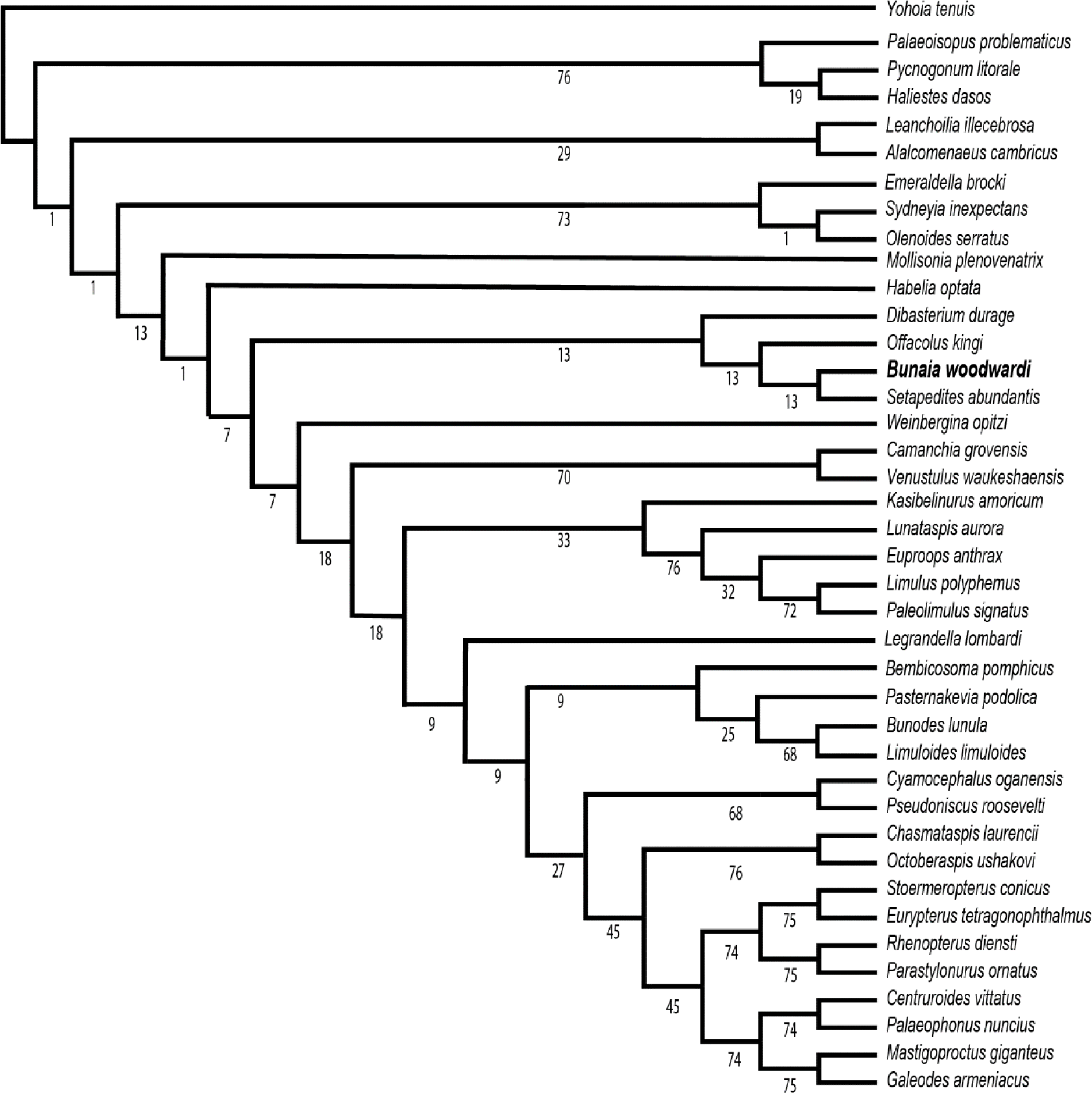
Phylogenetic position of *Bunaia woodwardi.* among euchelicerates. Parsimony phylogenetic analysis performed using random addition sequences followed by branch swapping with 100 000 repetitions, implied weight K= 6. Relative bremer support is shown with no brackets at the nodes. Matrix modified from Lamsdell [12] and Lustri *et al.* [8].

**Supplementary Figure 4.**
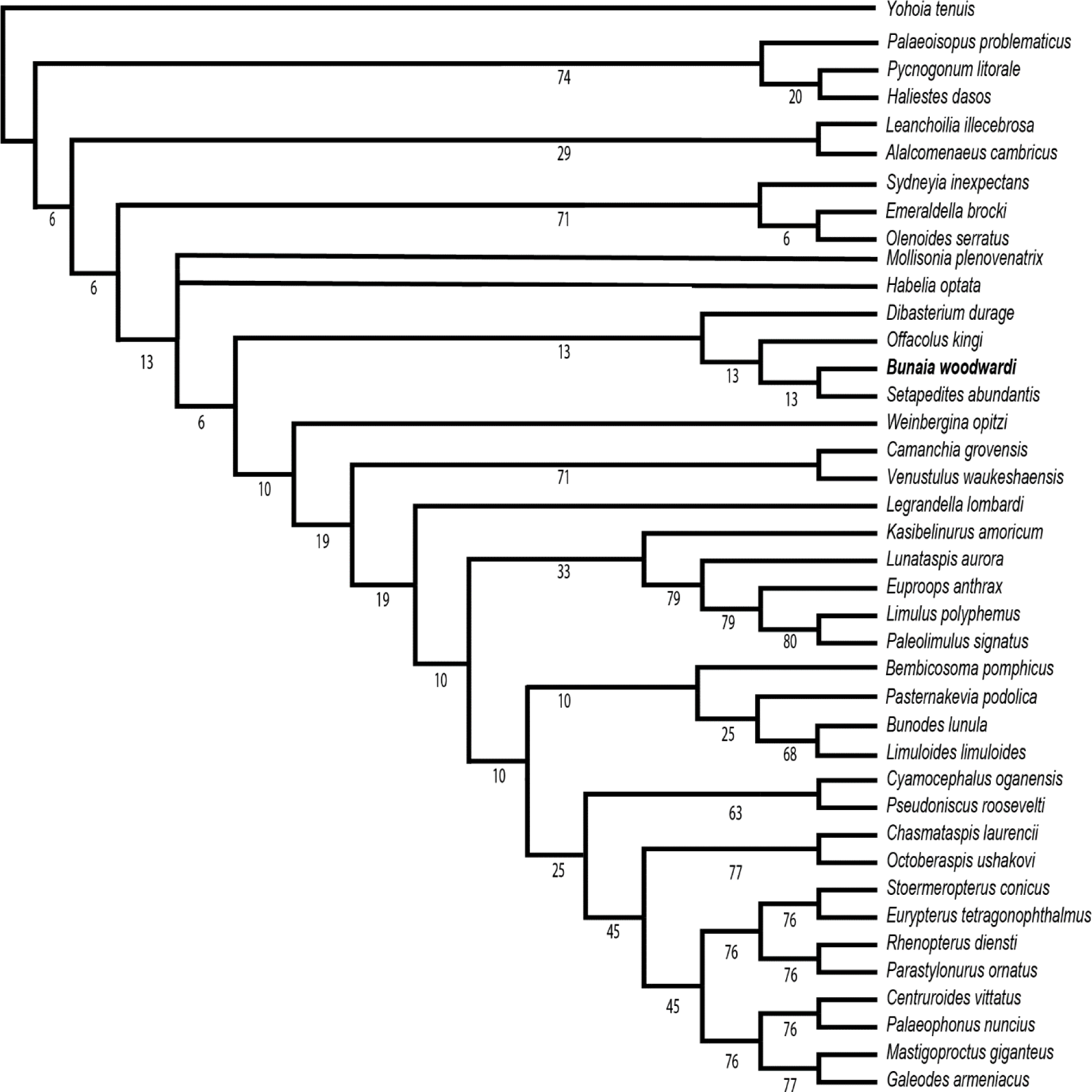
Phylogenetic position of *Bunaia woodwardi.* among euchelicerates. Parsimony phylogenetic analysis performed using random addition sequences followed by branch swapping with 100 000 repetitions, implied weight K= 9. Relative bremer support is shown with no brackets at the nodes. Matrix modified from Lamsdell [12].

## Notes

### Competing Interest Statement

The authors have declared no competing interest.

## References

1 Giribet, G. 2018 Current views on chelicerate phylogeny—a tribute to Peter Weygoldt. Zoologischer Anzeiger. 273, 7–13.

2 Giribet, G., Edgecombe, G. D. 2019 The phylogeny and evolutionary history of arthropods. Current Biology. 29, R592–R602.

3 Edgecombe, G. D. 2010 Arthropod phylogeny: an overview from the perspectives of morphology, molecular data and the fossil record. Arthropod Structure & Development. 39, 74–87.

4 Legg, D. A., Sutton, M. D., Edgecombe, G. D. 2013 Arthropod fossil data increase congruence of morphological and molecular phylogenies. Nature communications. 4, 1–7.

5 Wheeler, W. C., Hayashi, C. Y. 1998 The phylogeny of the extant chelicerate orders. Cladistics. 14, 173–192.

6 Dunlop, J., Arango, C. 2005 Pycnogonid affinities: a review. Journal of zoological systematics and evolutionary research. 43, 8–21.

7 Dunlop, J. A. 2010 Geological history and phylogeny of Chelicerata. Arthropod structure & development. 39, 124–142.

8 Lustri, L., Gueriau, P., Daley, A. in press Lower Ordovician synziphosurine reveals early euchelicerate diversity and evolution. Nature Communications.

9 Moore, R. A., McKENZIE, S. C., Lieberman, B. S. 2007 A Carboniferous synziphosurine (Xiphosura) from the Bear Gulch Limestone, Montana, USA. Palaeontology. 50, 1013–1019.

10 Zittel, K. v. Handbuch der Palaeontologie. I. Abtheilung, Palaeozoologie, 2 [Mollusca und Arthropoda]. R, Oldenbourg, München, Leipzig 1885.

11 Anderson, L. I., Selden, P. A. 1997 Opisthosomal fusion and phylogeny of Palaeozoic Xiphosura. Lethaia. 30, 19–31.

12 Lamsdell, J. C. 2013 Revised systematics of Palaeozoic ‘horseshoe crabs’ and the myth of monophyletic Xiphosura. Zoological Journal of the Linnean Society. 167, 1–27.

13 Briggs, D. E., Siveter, D. J., Siveter, D. J., Sutton, M. D., Garwood, R. J., Legg, D. 2012 Silurian horseshoe crab illuminates the evolution of arthropod limbs. Proceedings of the National Academy of Sciences. 109, 15702–15705.

14 Orr, P. J., Siveter, D. J., Briggs, D. E., Siveter, D. J., Sutton, M. D. 2000 A new arthropod from the Silurian Konservat–Lagerstätte of Herefordshire, UK. Proceedings of the Royal Society of London. Series B: Biological Sciences. 267, 1497–1504.

15 Sutton, M. D., Briggs, D. E., Siveter, D. J., Siveter, D. J., Orr, P. J. 2002 The arthropod Offacolus kingi (Chelicerata) from the Silurian of Herefordshire, England: computer based morphological reconstructions and phylogenetic affinities. Proceedings of the Royal Society of London. Series B: Biological Sciences. 269, 1195–1203.

16 Van Roy, P., Briggs, D. E., Gaines, R. R. 2015 The Fezouata fossils of Morocco; an extraordinary record of marine life in the Early Ordovician. Journal of the Geological Society. 172, 541–549.

17 Van Roy, P., Orr, P. J., Botting, J. P., Muir, L. A., Vinther, J., Lefebvre, B., El Hariri, K., Briggs, D. E. 2010 Ordovician faunas of Burgess Shale type. Nature. 465, 215–218.

18 Walcott, C. D. 1912 Cambrian Geology and Paleontology II: No 6--Middle Cambrian Branchiopdada, Malocrastraca, Trilobita, and Merostomata.

19 Aria, C., Caron, J.-B. 2017 Mandibulate convergence in an armoured Cambrian stem chelicerate. BMC evolutionary biology. 17, 1–20.

20 Simonetta, A. 1964 Osservazioni sugli Artropodi non Trilobiti della Burgess Shale (Cambriano Medio). III Contributo. I generi Molaria, Habelia, Emeraldella, Parahabelia (Nov.) Emeraldoides (Nov.). Monitore zool. ital. 72, 215–231.

21 Raasch, G. O. 1939 Cambrian merostomata. Geological Society of America.

22 Whittington, H. B. 1981 Rare arthropods from the Burgess Shale, Middle Cambrian, British Columbia. *Philosophical Transactions of the Royal Society of London. B*, Biological Sciences. 292, 329–357.

23 Ortega-Hernández, J., Legg, D. A., Braddy, S. J. 2013 The phylogeny of aglaspidid arthropods and the internal relationships within Artiopoda. Cladistics. 29, 15–45.

24 Clarke, J. M. 1919 I.—*Bunaia Woodwardi*, a New Merostome from the Silurian Waterlimes of New York. Geological Magazine. 6, 531–533.

25 Størmer, L. 1944 On the relationships and phylogeny of fossil and recent Arachnomorpha: a comparative study on Arachnida, Xiphosura, Eurypterida, Tribolita, and other fossil Arthropoda. Matematisk-Naturvidenskapelig Klasse. 5, 1–158.

26 Aria, C., Caron, J.-B. 2019 A middle Cambrian arthropod with chelicerae and proto-book gills. Nature. 573, 586–589.

27 Cotton, T. J., Braddy, S. J. 2003 The phylogeny of arachnomorph arthropods and the origin of the Chelicerata. Earth and Environmental Science Transactions of the Royal Society of Edinburgh. 94, 169–193.

28 Hou, X.-G. 1997 Arthropods of the lower Cambrian Chengjiang fauna, southwest China. Fossils and Strata. 45, 1–116.

29 Caron, J.-B., Rudkin, D. M., Milliken, S. 2004 A new late Silurian (Pridolian) naraoiid (Euarthropoda: Nektaspida) from the Bertie Formation of southern Ontario, Canada— delayed fallout from the Cambrian explosion. Journal of Paleontology. 78, 1138–1145.

30 Brett, C. E. 1998 Silurian-Early Devonian sequence stratigraphy, cycles and paleoenvironments of the Niagara Peninsula area of Ontario, Canada. Canada Geological Survey.

31 Ciurca Jr, S. Year Eurypterid biofacies of the Silurian–Devonian evaporite sequence: Niagara peninsula, Ontario, Canada and New York. New York State Geological Association 62nd Annual Meeting, Fredonia, New York; 1990; 1990. p. 1–30.

32 Ciurca Jr, S. J. Year Eurypterid horizons and the stratigraphy of Upper Silurian and Lower Devonian rocks of central-eastern New York State. New York State Geological Association 50th Annual Meeting, Syracuse, New York; 1978; 1978. p. 225–249.

33 Cramer, B. D., Brett, C. E., Melchin, M. J., Männik, P., Kleffner, M. A., McLaughlin, P. I., Loydell, D. K., Munnecke, A., Jeppsson, L., Corradini, C. 2011 Revised correlation of Silurian Provincial Series of North America with global and regional chronostratigraphic units and δ13Ccarb chemostratigraphy. Lethaia. 44, 185–202.

34 Lerosey-Aubril, R., Zhu, X., Ortega-Hernández, J. 2017 The Vicissicaudata revisited– insights from a new aglaspidid arthropod with caudal appendages from the Furongian of China. Scientific Reports. 7, 1–18.

35 Lerosey-Aubril, R., Ortega-Hernández, J. 2019 Appendicular anatomy of the artiopod Emeraldella brutoni from the middle Cambrian (Drumian) of western Utah. PeerJ. 7, e7945.

36 Brayard, A., Guériau, P., Thoury, M., Escarguel, G. 2019 Glow in the dark: Use of synchrotron μXRF trace elemental mapping and multispectral macro-imaging on fossils from the Paris Biota (Bear Lake County, Idaho, USA). Geobios. 54, 71–79.

37 Robin, N., Gueriau, P., Luque, J., Jarvis, D., Daley, A. C., Vonk, R. 2021 The oldest peracarid crustacean reveals a Late Devonian freshwater colonization by isopod relatives. Biology Letters. 17, 20210226.

38 Klug, C., Landman, N. H., Fuchs, D., Mapes, R. H., Pohle, A., Guériau, P., Reguer, S., Hoffmann, R. 2019 Anatomy and evolution of the first Coleoidea in the Carboniferous. Communications biology. 2, 1–12.

39 Lewis, P. O. 2001 A likelihood approach to estimating phylogeny from discrete morphological character data. Systematic biology. 50, 913–925.

40 Rudkin, D., Young, G. 2009 Horseshoe crabs–an ancient ancestry revealed. In Biology and conservation of horseshoe crabs. (ed. eds. pp. 25–44: Springer.

41 Størmer, L. 1934 Merostomata from the Downtonian sandstone of Ringerike, Norway. I Kommisjon hos Jacob Dybwad.

42 Eldredge, N., Smith, L. 1974 Revision of the suborder Synziphosurina (Chelicerata, Merostomata): with remarks on merostome phylogeny. American Museum novitates; no. 2543.

43 Nieszkowski, J. 1858 Zusätze zur Monographie der Trilobiten der Ostseeprovinzen: nebst der Beschreibung einiger neuen obersilurischen Crustaceen. Oxford Microform Publications.

44 Woodward, H. 1868 I.—On a New Limuloid Crustacean [Neolimulus falcatus] from the Upper Silurian of Lesmahagow, Lanarkshire. Geological Magazine. 5, 1–3.

45 Clarke, J. M. 1902 Notes on Paleozoic crustaceans. University of the State of New York.

46 Ruedemann, R. 1916 Account of some new or little-known species of fossils, mostly from the Paleozoic rocks of New York. New York State Museum Bulletin. 189, 7–112.

47 Anderson, L. I. 1999 A new specimen of the Silurian synziphosurine arthropod Cyamocephalus. Proceedings of the Geologists Association. 110, 211–216.

48 Moore, R. A., Briggs, D. E., Braddy, S. J., Anderson, L. I., Mikulic, D. G., Kluessendorf, J. 2005 A new synziphosurine (Chelicerata: Xiphosura) from the Late Llandovery (Silurian) Waukesha Lagerstatte, Wisconsin, USA. Journal of Paleontology. 79, 242–250.

49 Stürmer, W., Bergström, J. 1981 Weinbergina, a xiphosuran arthropod from the Devonian Hunsrück Slate. Paläontologische Zeitschrift. 55, 237–255.

50 Ortega-Hernández, J., Janssen, R., Budd, G. E. 2017 Origin and evolution of the panarthropod head–a palaeobiological and developmental perspective. Arthropod structure & development. 46, 354–379.

51 Budd, G. E. 2021 The origin and evolution of the euarthropod labrum. Arthropod structure & development. 62, 101048.

52 Haug, J. T., Briggs, D. E., Haug, C. 2012 Morphology and function in the Cambrian Burgess Shale megacheiran arthropod Leanchoilia superlata and the application of a descriptive matrix. BMC Evolutionary Biology. 12, 1–20.

53 Brenneis, G., Ungerer, P., Scholtz, G. 2008 The chelifores of sea spiders (Arthropoda, Pycnogonida) are the appendages of the deutocerebral segment. Evolution & development. 10, 717–724.

54 Bruton, D. L. 1981 The arthropod Sidneyia inexpectans, Middle Cambrian, Burgess Shale, British Columbia. Philosophical Transactions of the Royal Society of London. B, Biological Sciences. 295, 619–653.

55 Stein, M. 2013 Cephalic and appendage morphology of the Cambrian arthropod Sidneyia inexpectans. Zoologischer Anzeiger-A Journal of Comparative Zoology. 253, 164–178.

56 Walcott, C. D. 1911 Cambrian Geology and Paleontology II: No. 2--Middle Cambrian Merostomata.

57 Stein, M., Selden, P. A. 2012 A restudy of the Burgess Shale (Cambrian) arthropod Emeraldella brocki and reassessment of its affinities. Journal of Systematic Palaeontology. 10, 361–383.

58 Walcott, C. D. 1912 Middle Cambrian Branchiopoda, Malacostraca, Trilobita, and Merostomata. Smithsonian Institution.

59 Lerosey-Aubril, R., Ortega-Hernandez, J., Zhu, X. 2013 The first aglaspidid sensu stricto from the Cambrian of China (Sandu Formation, Guangxi). Geological Magazine. 150, 565–571.

60 Whittington, H., Chatterton, B., Speyer, S., Fortey, R., Owens, R., Chang, W., Dean, W., Jell, P., Laurie, J., Palmer, A. 1997 Treatise on Invertebrate Paleontology, Part O, Arthropoda 1, Trilobita, Revised. Geological Society of America, Boulder, CO and University of Kansas, Lawrence, KS. 530.

61 Ewington, D., Clarke, M., Banks, M. Year A Late Permian fossil horseshoe crab (Paleolimulus: Xiphosura) from Poatina, Great Western Tiers, Tasmania. Papers and Proceedings of the Royal Society of Tasmania; 1989; 1989. p. 127–131.

62 Lerosey-Aubril, R., Paterson, J. R., Gibb, S., Chatterton, B. D. 2017 Exceptionally-preserved late Cambrian fossils from the McKay Group (British Columbia, Canada) and the evolution of tagmosis in aglaspidid arthropods. Gondwana Research. 42, 264–279.

63 Edgecombe, G. D., García-Bellido, D. C., Paterson, J. R. 2011 A new leanchoiliid megacheiran arthropod from the lower Cambrian Emu Bay Shale, South Australia. Acta Palaeontologica Polonica. 56, 385–400.

64 Briggs, D. E., Siveter, D. J., Siveter, D. J., Sutton, M. D., Legg, D., Lamsdell, J. C. 2023 A vicissicaudatan arthropod from the Silurian Herefordshire Lagerstätte, UK. Royal Society Open Science. 10, 230661.

65 Stein, M., Church, S. B., Robison, R. A. 2011 A new Cambrian arthropod, Emeraldella brutoni, from Utah. Paleontological Contributions. 2011, 1–9.

66 Broili, F. 1932 Eine neue Crustacee aus dem rheinischen Unterdevon.

67 Chlupác, I. 1988 The enigmatic arthropod Duslia from the Ordovician of Czechoslovakia. Palaeontology. 31, 611–620.

68 Riedl, R. 1978 Order in living organisms: a systems analysis of evolution. John Wiley & Sons.

69 Van Roy, P. 2005 An aglaspidid arthropod from the Upper Ordovician of Morocco with remarks on the affinities and limitations of Aglaspidida. Earth and Environmental Science Transactions of the Royal Society of Edinburgh. 96, 327–350.

70 Van Roy, P. 2006 Non-trilobite arthropods from the Ordovician of Morocco: Ghent University.

71 Patterson, C. 1982 Morphological characters and homology. Problems of phylogenetic reconstruction. 21–74.

72 Hesselbo, S. P. 1992 Aglaspidida (Arthropoda) from the upper Cambrian of Wisconsin. Journal of Paleontology. 885–923.

